# Discovery of a Well-Folded Protein Interaction Hub Within the Human Long Non-Coding RNA *NORAD*

**DOI:** 10.1101/2023.08.07.552337

**Authors:** Ananthanarayanan Kumar, Han Wan, Zion Perry, Shivali Patel, Rafael Tavares, Anna Marie Pyle

## Abstract

The long non-coding RNA *NORAD* functions in maintaining genomic stability in humans via sequestering Pumilio proteins from the cytoplasm, and thereby modulating the gene expression of mRNA targets of Pumilio proteins. Despite its role in fundamental cellular pathways including chromosome segregation and DNA damage response, there have been limited structural and biophysical descriptions of *NORAD*. Here, using an integrative approach combining chemical probing coupled to high throughput sequencing, and RNA-pull downs coupled with mass spectrometry, we discovered a well-folded and structured protein interaction hub within the functional core of *NORAD*. Our *in vitro* biochemical reconstitutions using purified recombinant proteins and a *NORAD* repeat unit region within this hub reveal the assembly of a higher-order multimeric RNA-protein complex.

## INTRODUCTION

A majority of the mammalian genome is transcribed into large non-coding RNAs (>500nt) that do not have any protein-coding ability, and such non-coding transcripts have been implicated in regulating a plethora of key cellular properties such as chromosome accessibility, X-chromosome inactivation, R-loop formation and gene expression regulation (Cech and Steitz 2014, Yao, Wang et al. 2019, Mattick, Amaral et al. 2023).) (Mattick, Amaral et al. 2023). With the advent of high throughput sequencing technology and genome engineering methods, we are beginning to discover several key roles played by lncRNAs in fundamental cellular pathways (Kashi, Henderson et al. 2016, Yao, Wang et al. 2019, Haswell, Mattioli et al. 2021). *NORAD* (Non-Coding RNA Activated By DNA Damage) is one such lncRNA in humans that is ∼ 5.5 kb long, abundantly expressed and highly conserved among mammals (Lee, Kopp et al. 2016, Tichon, Gil et al. 2016, Munschauer, Nguyen et al. 2018, Stojic, Lun et al. 2020). *NORAD* was discovered in a seminal publication by the Mendel lab while studying lncRNAs involved in DNA damage response (Lee, Kopp et al. 2016). The study thoroughly characterized the function of *NORAD* in maintaining genomic stability in the cells by means of its interaction with Pumilio proteins. Simultaneously, studies from the Ulitsky lab also identified *NORAD* as a key player in chromosome segregation via its interaction with Pumilio proteins (Tichon, Gil et al. 2016). Furthermore *NORAD* knock-down cells displayed reduced DNA replication fork velocity and defects in chromosome segregation (Munschauer, Nguyen et al. 2018, Elguindy, Kopp et al. 2019).

*NORAD* has been shown to be involved in genome stability maintenance via its interaction with the Pumilio proteins and formation of phase separated condensates in the cytoplasm (Elguindy and Mendell 2021). Human Pumilio proteins belong to the PUF family of sequence specific RNA-binders that post-transcriptionally control messenger RNA fate (mRNA) (Goldstrohm, Seay et al. 2007, Wolfe, Schagat et al. 2020, Enwerem, Elrod et al. 2021). *NORAD* contains at least seventeen Pumilio response elements (PREs) throughout its sequence **(Figure 1B)** (Tichon, Gil et al. 2016). PREs are 8-nt long 5′-*UGUAHAUA*−3′ consensus binding sites for mammalian Pumilio proteins (Galgano, Forrer et al. 2008, Goldstrohm, Hall et al. 2018). *NORAD* functions by sequestering RNA-binding Pumilio proteins via its PREs in a regulated manner, forming phase separated condensates, thereby preventing the targeted degradation of cognate mRNAs, which are involved in cell division, DNA replication, and the DNA damage response (Elguindy and Mendell 2021).

**Figure 1:**
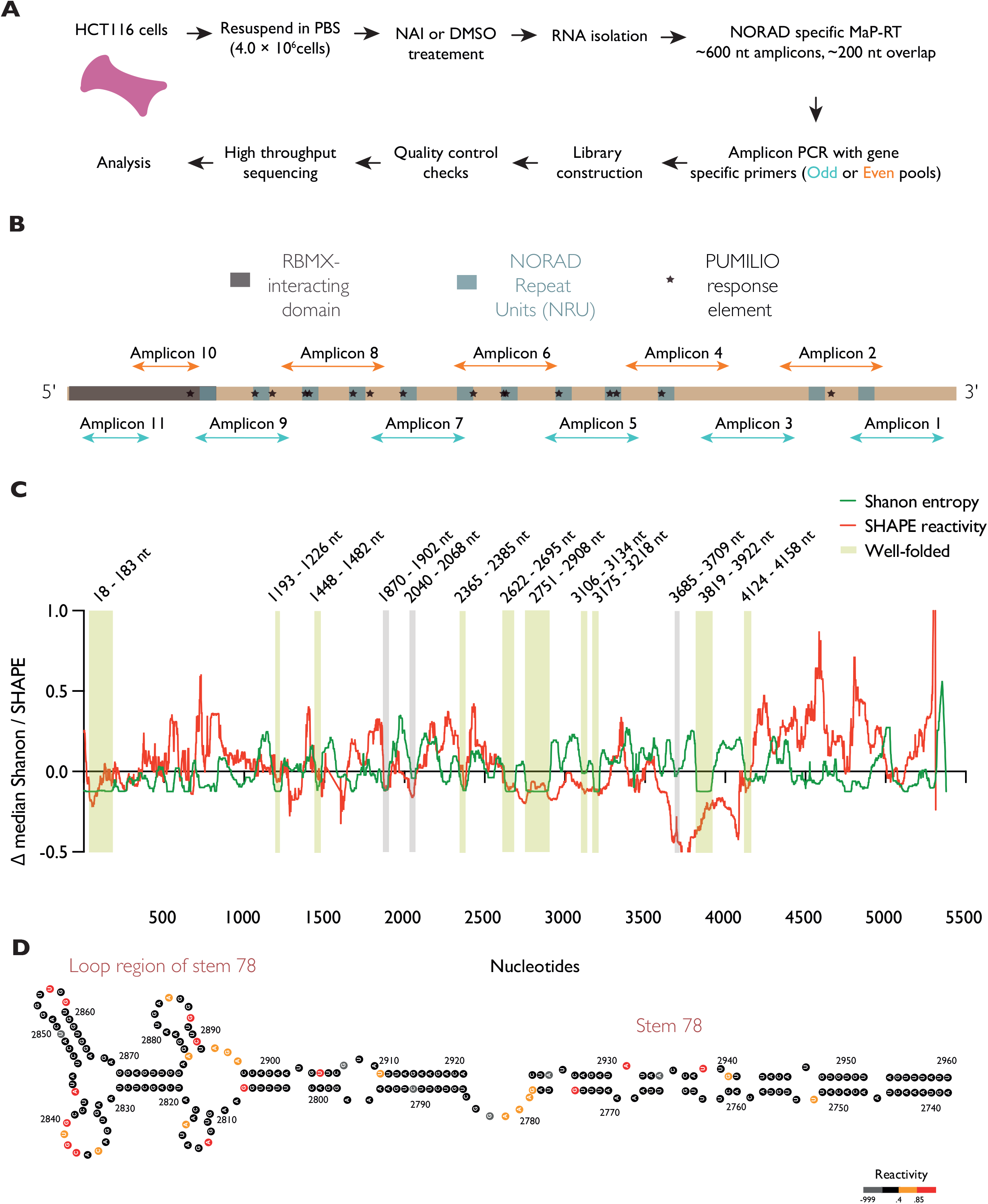
*In cell* SHAPE-MaP of *NORAD* reveals several well-folded structural motifs. **A:** Schematic of the *in cell* SHAPE-MaP method used in the study. **B:** Domain diagram of *NORAD* showing all repeat units, previously identified protein binding sites, and SHAPE-MaP amplicon design. **C:** Analysis of Shannon entropy and SHAPE reactivities calculated from *in cell* SHAPE-MaP experiment reveals > 10 highly structured, well-determined domains within *NORAD* (data from Rep3). Yellow boxes represent well-folded motifs found in both the replicates. Gray boxes represent well-folded domains found in only one of the two replicates. **D:** Minimum free energy secondary structure model of the 78 stem found between *NORAD* repeat units 7 & 8 (*NRU78*). The structures were visualized using the program StructureEditor.

Studies have shown that lncRNAs form complex secondary and tertiary structures, and long-range interactions can mediate their cellular function (Novikova, Hennelly et al. 2013, Hawkes, Hennelly et al. 2016, Uroda, Anastasakou et al. 2019). Due to their large size and difficulty in purifying a native-folded state, there are only a handful of cases of an experimentally determined *in vitro* secondary structural profile of a lncRNA (Novikova, Hennelly et al. 2012, Ilik, Quinn et al. 2013, Somarowthu, Legiewicz et al. 2015, Hawkes, Hennelly et al. 2016, Liu, Somarowthu et al. 2017, Jones and Sattler 2019, Uroda, Anastasakou et al. 2019). When it comes to probing these structures within cells, there are significant challenges such as lack of abundance, which leads to difficulty in amplification and presence of alternate isoforms, leading to heterogeneity in structure (Smola and Weeks 2018, Busan, Weidmann et al. 2019). In the case of *NORAD*, sequence analysis revealed the presence of 12 repeated structural units called as *NORAD* repeat units (*NRU*s) **(Figure 1B)** (Tichon, Gil et al. 2016, Tichon, Perry et al. 2018). Many of the *NRU*s contain one or two PREs followed by a short stem, a stretch of U-rich sequences, and a longer stem followed by an A/G-rich sequence (Tichon, Perry et al. 2018). Despite the presence of predicted structural features within *NORAD*, and their role in fundamental cellular pathways, there are limited biophysical descriptions of *NORAD* RNA. The lack of an in-depth secondary structure map further impedes any detailed studies of *NORAD* and its mechanism of maintaining genome stability. Using the nextPARS method on *in vitro* produced *NORAD* truncations, a recent study characterized the repeat unit architecture present within the RNA (Chorostecki, Saus et al. 2021). A recent preprint from the Ulitsky lab reported long-range interactions within *NORAD* in cells and elaborated on the structural domain organization (Ziv, Farberov et al. 2021). To further contribute to this growing interest in understanding RNA structures within *NORAD*, we employed the SHAPE-MaP pipeline to determine the complete secondary structure of human *NORAD* at single-nucleotide resolution in HCT116 cells (Smola and Weeks 2018). Our comprehensive structural probing revealed a network of well-folded stable RNA structures within *NORAD,* including the regions between nucleotides 14-190 (RBMX-binding site) and between 2620-2900 (within a previously identified minimal functional unit of *NORAD*). Previous proteomic studies have found that *NORAD* interacts with a plethora of accessory proteins (other than Pumilio) involved in nucleic acid processing and DNA damage response (Munschauer, Nguyen et al. 2018, Spiniello, Knoener et al. 2018, Tichon, Perry et al. 2018). To understand where these proteins might assemble on *NORAD*, we probed the RNA in cellular extracts depleted of proteins and compared this to the *in cell* SHAPE data, leading to the discovery of five potential protein binding hubs, all of which were selected in an unbiased manner. Surprisingly, one of these newly discovered protein hotspots lies within the *NORAD* repeat units 7 & 8 (*NRU78*), which was also found to be in a highly structured and well-folded region of the RNA (low-Shannon and low-SHAPE metric, hereafter referred to as lowSS) (Weeks 2021). Next, using *in vitro* RNA-pull-downs coupled with a mass spectrometry, we identified well-known binders of *NORAD,* such as Pumilio1 and FUBP3, and discovered several novel protein binding partners of *NORAD NRU78* including HARS2 and CELF1. Finally, using purified Pumilio1 and FUBP3 proteins, and a construct comprised of *NRU7*, we reconstituted a minimal *NORAD* RNA-protein complex (RNP). Taken together, the chemical probing and proteomics data have enabled the identification of a well-folded protein interaction hub within *NORAD*.

## RESULTS

### In Cell SHAPE-MaP of *NORAD* Reveals Well-Folded RNA Structural Motifs

To determine the secondary structure of *NORAD*, the full-length RNA was probed in HCT 116 cells using the SHAPE-MaP strategy (Smola, Rice et al. 2015, Smola and Weeks 2018) **(Figure 1A)**. 48h post-seeding, HCT116 cells were treated with either the electrophilic SHAPE reagent NAI or control DMSO. NAI modifies 2′-OH on flexible RNA nucleotides, so single stranded and flexible regions in the RNA tend to have higher SHAPE reactivities than double stranded or protein/nucleic acid bound regions in the cell (Smola and Weeks 2018, Velema and Kool 2020). Next, the RNA was extracted from SHAPE-probed cells, and sequencing libraries were generated using the tiled amplicon approach (Huston, Wan et al. 2021). NAI-modified nucleotides in *NORAD* were recorded as internal mutations during reverse transcription using gene specific primers and were subjected to next generation sequencing (Siegfried, Busan et al. 2014). Two replicates were performed, and the resulting sequencing data was used to generate a SHAPE reactivity profile for *NORAD* using the ShapeMapper2 pipeline (Busan and Weeks 2018). Effective reactivity data were collected for >95% of nucleotides in both replicates. Analyzing the mutational rates of the NAI-treated versus the control DMSO-treated sample revealed a significant increase in the mutational rates of the former, confirming that the full-length *NORAD* was successfully modified with NAI *in vivo* **(Supplemental Figure 1A)**. Comparison of the normalized SHAPE reactivities of the two replicates revealed a high correlation between the data sets (Pearson’s r = 0.94 & Spearman’s r = 0.81) **(Supplemental Figure 1B)**. Both the replicates also successfully passed the SHAPEmapper quality control checks (**Supplemental Figure 1C).** Next, the secondary structure of full-length *NORAD* and the Shannon entropy values per nucleotide were determined using the SuperFold pipeline with the experimentally determined SHAPE reactivities as constraints (Smola, Rice et al. 2015). Shannon entropy is a measure of base-pair probabilities across all possible conformations during structure prediction (Garcia-Martin and Clote 2015). Visualizing the full-length secondary structure of *NORAD* with the SHAPE reactivity superimposed on the corresponding nucleotides revealed that the overall reactivities match well with the predicted structures i.e. structured regions have low SHAPE values, and the loop regions have medium or high SHAPE values. Using the low-SHAPE and low-Shannon metric (lowSS) (Weeks 2021), ten well-folded domains were identified in *NORAD* **(Figure 1C)**. These regions were identified in both the replicates (**Supplement Figure 2).** Several of these domains lie within regions in *NORAD* previously attributed to protein binding. Some of these include the regions between 14-190 nt of *NORAD* (**Supplement Figure 3**) implicated in binding the protein RBMX. Other regions of interest include a well-folded region between 1193-1226 nts of *NORAD*. This falls in the 3′-end of the *NORAD* repeat unit 2 that harbors a SAM-68 binding site (Tichon, Perry et al. 2018). Regions 2342-2407 nts are also found to be well-folded from the lowSS analysis. Notably, this region consists of the beginning of the *NORAD* repeat unit 6 (*NRU6*) and extends to the end of its ‘long stem’ (**Supplemental Figure 3**). The repeat unit 7 of *NORAD* (*NRU7*) consists of two PREs, a characteristic short stem followed by a long stem and lies within one of the well-folded domain (2609-2704 nt) discovered in this study. Interestingly, the lowSS regions lying within nucleotides 2749-2915 of *NORAD* is found immediately after the *NRU 7*, and includes a ∼ 200 nt long stem (referred to as 78 Stem hereafter) **(Figure 1D, Supplemental Figure 3)**. This well-folded stem is situated in between the repeat units *NRU7* and *NRU8*. Similarly, between the repeat units *NRU8* and *NRU9*, there is a well-folded short structure between nucleotides 3103-3132. The regions spanning nucleotides 3809-3935 of *NORAD* also consist of many well-folded structures (**Supplement Figure 3**). This domain has not yet been implicated in any *NORAD* function and it does not lie within any repeat unit structures. Notably, the 3′-end of the *NORAD* broadly consists of high SHAPE reactivity regions and contains many large loops **(Figure 1D, Supplemental Figure 3)**.

### Unbiased Identification of Protein Binding Hubs in *NORAD* using Chemical Probing

To explore how the cellular environment influences the structure of *NORAD*, and to discover protein binding sites, we probed *NORAD* in HCT116 cell extracts depleted of proteins (Smola, Calabrese et al. 2015, Smola, Christy et al. 2016). HCT116 cells were gently lysed open, followed by incubation with a proteolytic enzyme Proteinase K (Patel, Sexton et al. 2023) **(Figure 2A)**. This ensures the removal of all cognate protein partners of *NORAD* RNA present in the lysate. The proteolyzed cellular extract is then treated with NAI or DMSO, and subsequent RNA purification, SHAPE-library preparation, sequencing and data analysis were done similar to the *in cell* experiment described in the previous section. The experiment was done in replicates. The effective reactivity data was collected for >95% of nucleotides in both replicates. The NAI-treated samples had significantly higher mutation rates in comparison to the control DMSO-treated sample, confirming that the full-length *NORAD* was successfully modified with NAI in the cell extracts (*ex cellulo*) **(Supplemental Figure 4A)**. Comparison of the normalized SHAPE reactivities of the two replicates revealed a high correlation between the data sets (Pearson’s r = 0.7861 & Spearman’s r = 0.8468) **(Supplemental Figure 4B)**. Sequencing data from both the replicates also successfully passed the SHAPEmapper quality control checks (**Supplemental Figure 4C).** Next, the secondary structure of full-length *NORAD* and the Shannon entropy values per nucleotide were calculated using the SuperFold pipeline with the *ex cellulo* SHAPE reactivities as constraints. Similar to the previous analysis, the lowSS metric was used to pinpoint well-folded domains in *NORAD* **(Supplement Figure 5)**. In total, 16 such regions were identified, in both replicates **(Supplement Figure 5).**

**Figure 2:**
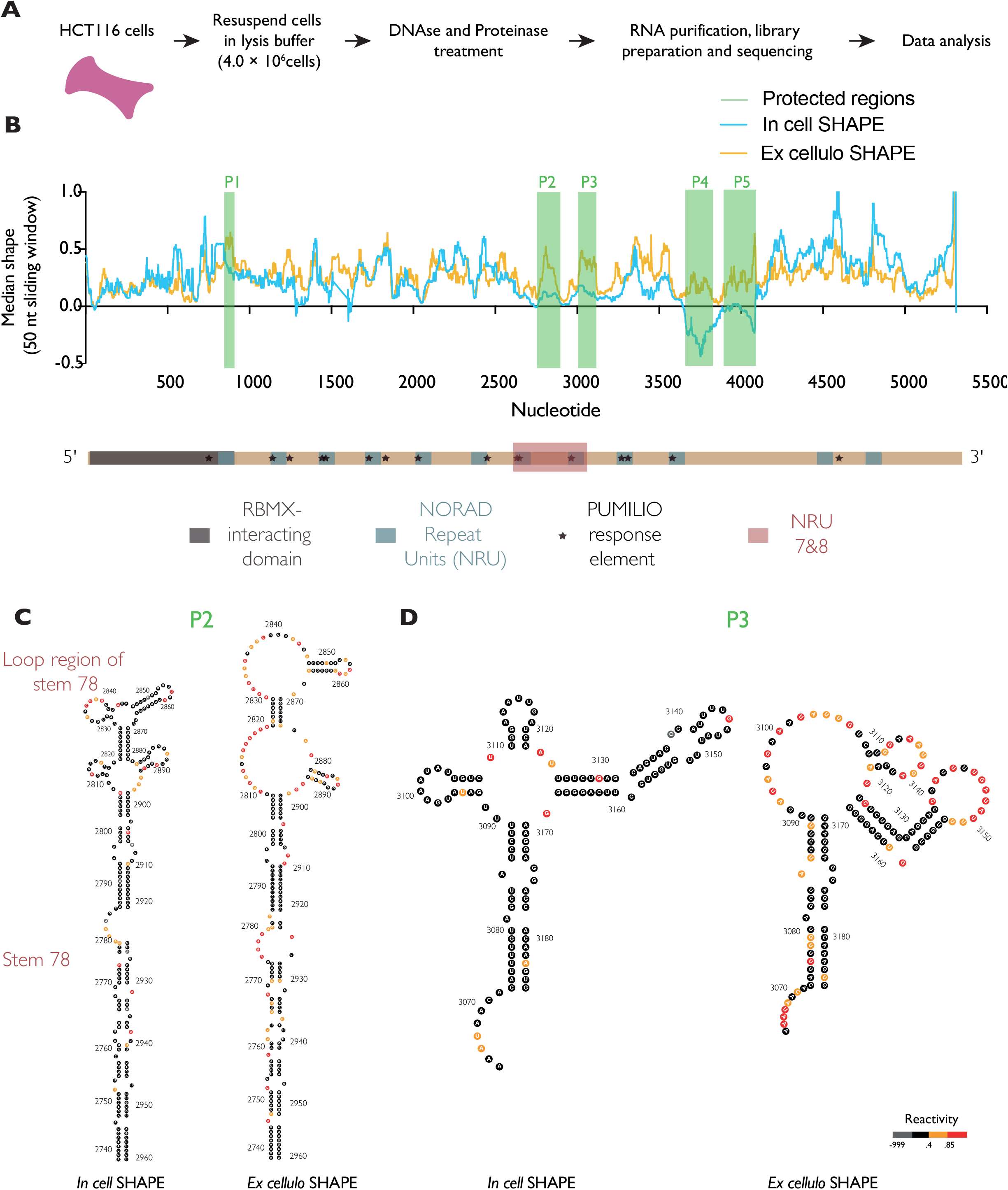
Identification of protein binding hubs within *NORAD* using *ex cellulo* SHAPE-MaP. **A:** Schematic of the *ex cellulo* SHAPE-MaP method used in the study. **B:** Comparison of *in cell* and *ex cellulo* SHAPE reactivities in 50-nt sliding windows reveals potential protein binding hubs within *NORAD*. Green boxes represent regions with the highest difference between *in cell* and *ex cellulo* SHAPE reactivity. Two strong potential protein binding hubs are identified within *NORAD* repeat units 7 and 8 (*NRU78*). **C:** Comparison of *in cell* and *ex cellulo* SHAPE reactivities of *NRU7&8* superimposed on the predicted secondary structure profiles. **D:** Comparison of *in cell* and *ex cellulo* SHAPE reactivities of a region immediately downstream of *NORAD* repeat unit 8 (*NRU8)* superimposed on the predicted secondary structure profiles.

Next, by comparing the SHAPE reactivity changes between the *in cell* and *ex cellulo* experiments, and analyzing the predicted secondary structure model, we identified five potential protein binding hubs within *NORAD* **(Figure 2B)**. These five hubs (referred to as P1 - P5) were identified in all the replicates and across all four samples that were compared with each other (**Supplemental Figure 6).** P1 overlaps with the *NORAD* repeat unit 1 (*NRU1*) and is neither well-folded nor stable according to the lowSS criteria, thereby suggesting that *NRU 1* is likely a dynamic protein binding hub (**Supplemental Figure 7)**. Interestingly, P2 corresponds to the 78 Stem that lies between *NRU 7* and *NRU 8*, which was previously identified in the *in cell* SHAPE experiment **(Figure 2B)**. P3 lies immediately after *NRU8* and P4 and P5 lies in an uncharacterized part of *NORAD* between *NRU 10* and *NRU 11* **(Figure 2B)**. The Shannon entropy and SHAPE reactivity profiles of the *in cell* and *ex cellulo* samples revealed that the well-folded domains in *NORAD* overlap between these samples at the following regions: the 5′-domain containing the RBMX binding site, the domain encompassing NRU7 and NRU8 (hereafter referred to as NRU7&8) and regions 3806 - 3930 nt after *NRU10* (**Supplemental Figure 7)**. Furthermore, comparing the lowSS regions between the *in cell* and *ex cellulo* samples highlights that the protein binding hub P2 is well-folded in cells, and becomes highly reactive and flexible upon removal of proteins (**Supplemental Figure 7)**. This change in reactivity is a result of a structural change on the 78 stem in the absence of proteins and can be visualized in the predicted secondary structure of this region **(Figure 2C**). Similarly, the P3 region undergoes structural rearrangements upon removal of proteins **(Figure 2C**).

A region of particular interest is the domain *NRU78* since it harbors two well-folded repeat units, interspersed by an unusually long 200 nucleotides stem that also serves as a protein binding hotspot (P2). Together, it can be inferred that this domain is a highly structured protein binding hub within *NORAD*. Interestingly, previous studies have shown that the region spanning nucleotides 2494-3156 of *NORAD*, which contains *NRU78* (referred to as ND4 in their study), is highly conserved in mammals, contains multiple binding sites for Pumilio proteins and is sufficient to rescue the chromosome segregation phenotype of HCT116 cells lacking *NORAD* (Elguindy, Kopp et al. 2019, Elguindy and Mendell 2021). To better understand the molecular details of how *NORAD* functions, it is imperative to identify the proteome associated with this ‘core’ region of *NORAD* and to characterize its molecular architecture.

### Discovery of Known and Novel Cognate Protein Partners of *NORAD*

There have been several published studies of proteomics characterization of *NORAD* (Munschauer, Nguyen et al. 2018, Spiniello, Knoener et al. 2018, Tichon, Perry et al. 2018). These studies either used *in vitro* produced RNA purified under denaturing conditions, or they were performed on full-length RNA in the cellular context. As a result, despite knowing the list of proteins that associate with the full-length RNA, the proteome associated with natively folded individual *NORAD* domains remains unknown. The current report, and several recent studies have highlighted the importance and abundance of RNA structure in *NORAD* (Chorostecki, Saus et al. 2021, Ziv, Farberov et al. 2021). Therefore, it is important to purify *NORAD* RNA domains under non-denaturing conditions for any biophysical or chemical studies. To identify the protein binding partners of specific domains in *NORAD*, we devised a strategy consisting of i*n vitro* RNA-pull downs coupled with label-free quantitative mass spectrometry, using purified RNA under native conditions (Ilagan and Jurica 2014, Chillon, Marcia et al. 2015, Di Tomasso, Miller Jenkins et al. 2016) (**Figure 3A**).

**Figure 3:**
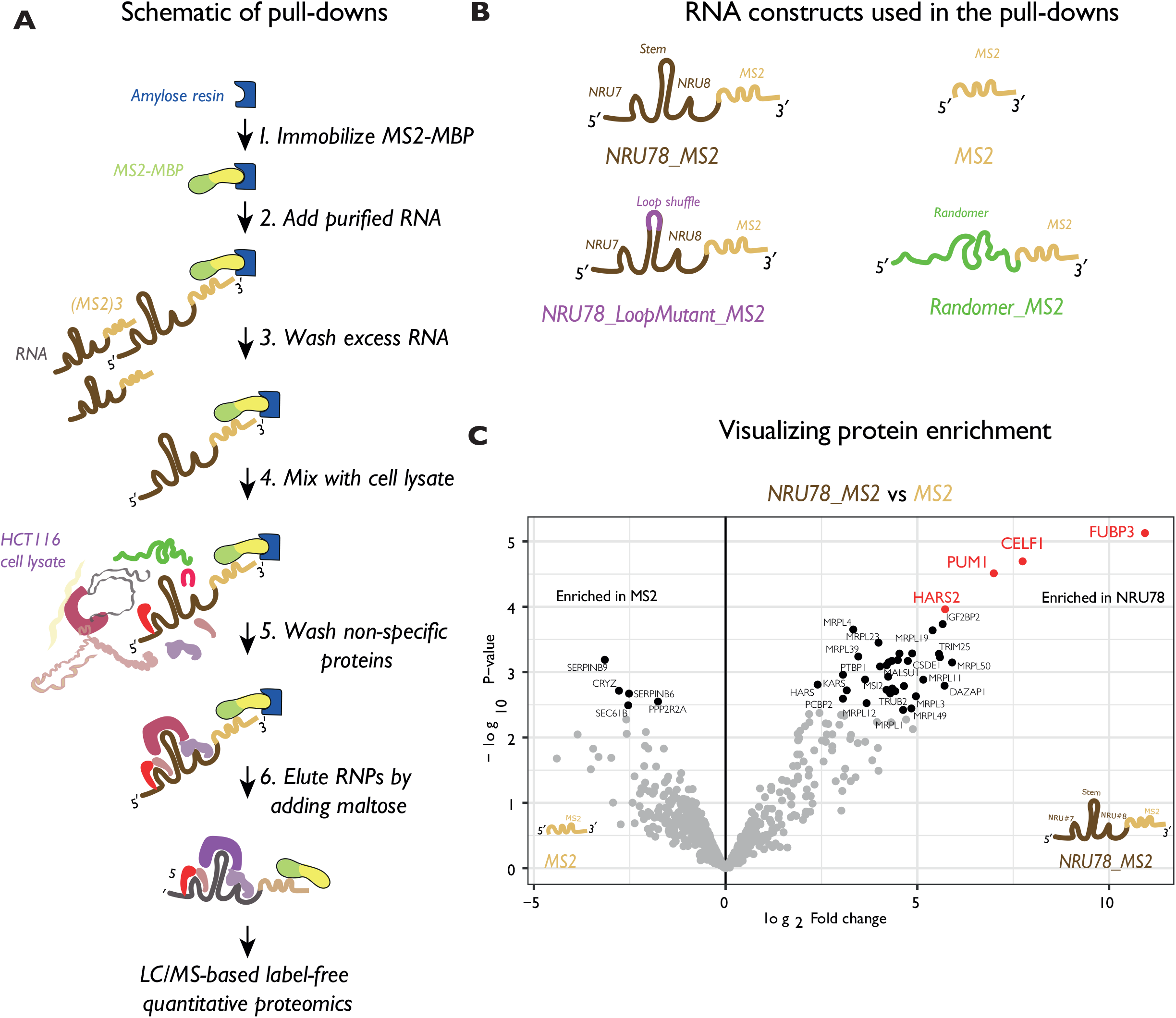
*In vitro* pull-downs coupled with quantitative mass spectrometry identified protein partners of *NORAD NRU78* **A:** Schematic of the *MS2*-lncRNA pull-downs used in the study. **B:** Cartoon representation of the RNA constructs used in the study. *NRU78_MS2* consists of the repeat units 7 & 8 of *NORAD* fused to a triple *MS2* stem in its 3′-end. The *NRU78_LoopMutant_MS2* consists of the repeat units 7 & 8 of *NORAD* with its *78 stem loop* mutated with a shuffled sequence, and fused to a triple *MS2* sequence in its 3′-end. *MS2* consists of a triple *MS2* stem loop sequence. *Randomer_MS2* consists of a randomized sequence with the same base composition and size as *NORAD* repeat units 7 & 8, and fused to a triple *MS2* stem in its 3′-end. **C:** Volcano plots showing specific enrichment of proteins in *NORAD NRU78* pull-downs in comparison with *MS2* alone control pull-downs.

Here, we focused on *NRU78* that lies within the functional core of *NORAD*, with the aim of identifying accessory protein partners in addition to known binders such as Pumilio. First, an RNA construct consisting of *NRU78* was fused to a triple *MS2* stem loop at its 3′-end. The *MS2* stem is used as a handle to immobilize *NORAD* domain *NRU78* onto amylose resins that are saturated with *MBP-MS2* proteins. The RNA loaded resins are used as a bait to isolate and purify *NORAD* specific protein complexes by incubating with HCT116 cellular extracts (**Figure 3A**). As controls, the following RNA constructs were used: 1) triple *MS2* stem loop alone, 2) a randomized sequence of *NRU78* fused to a triple *MS2* stem loop and 3) *NRU78* with a mutated loop in the 78 Stem, fused to a triple *MS2* stem loop (**Figure 3B**). The ‘randomer’ and ‘*MS2* alone’ controls are to rule out nonspecific interactors from *NORAD* specific proteins. The construct containing a mutation in the loop of 78 Stem serves as an additional control, where proteins binding to the 78 Stem loop will not be isolated anymore. The four RNA constructs were *in vitro* transcribed and purified using the native purification method (Chillon, Marcia et al. 2015). The purity and homogeneity of the RNAs were assessed by size exclusion chromatography and denaturing Urea-PAGE (**Supplemental Figure 8A and 8B**). The specific RNPs isolated in the pull-down experiments were run on a SDS-PAGE to visualize the protein content, and denaturing Urea-PAGE to ensure that the eluted RNPs still contained intact RNA at the correct size (**Supplemental Figure 8B and 8C**). To ensure that proteins specific to *NORAD* were being isolated, the elution samples were analyzed by Western blot using a polyclonal antibody specific to human Pumilio1 (**Supplemental Figure 8D**). The pull-downs and subsequent liquid chromatography with tandem mass spectrometry (LC-MS-MS) analysis were done in replicates. A total of ∼ 600 proteins were identified per each sample. Differential Enrichment analysis of Proteomics data (DEP) analysis was performed to identify specifically enriched proteins in different samples. A correlation matrix is used to visualize the Pearson correlation between the normalized lable-free quantification (LFQ) intensity across the replicates (**Supplemental Figure 9A**). A heatmap representation of all significant proteins in all the eight samples shows a reasonable clustering of proteins across replicates and within each sample (**Supplemental Figure 9B**). The plot further provides a global overview of how proteins enriched in the *NORAD NRU78* samples are not well represented in the ‘randomer’ and ‘*MS2* alone’ control samples.. In order to inspect the specific enrichment of proteins in *NORAD NRU78* pull downs, we defined high confidence interactomes as log_2_fold change >1.5 and p value < 0.05. Comparing *NORAD NRU78* pull downs to *MS2* alone controls, we discovered several known binding partners of *NORAD* such as Pumilio1 and FUBP3 (**Figure 3C**). In addition, several novel *NRU78* associated proteins were discovered and these include the RRM-domain containing CELF1 protein, aminoacyl-tRNA synthetases HARS2 and others. Reassuringly, several of these interactors including PUM1, FUBP3, CELF1 and HARS2 were also found to be highly enriched in the comparison between *NRU78* and the randomer sample (**Supplemental Figure 9C**). The full list of *NORAD NRU78* interactors is presented in **Supplementary Table 1**. Next, to gain insight into the factors that specifically bind to the loop region in the 78 stem, the enrichment of proteins between *NORAD NRU78* and *NORAD NRU78* loop mutant were plotted (**Supplemental Figure 9D**). HARS2 was found to be at the top in the list of enriched proteins. HARS1 was also additionally found to be enriched in this analysis. Surprisingly, several mitochondrial ribosomal subunits such as MALSU1, MRPL3 and MRPL18 were found to be associated with the 78 stem loop.

### Biochemical Reconstitution of a Minimal *NORAD*-Pumilio-FUBP3 Complex

To confirm the direct binding of the newly identified factors with *NORAD*, we used an *in vitro* reconstituted system using purified recombinant proteins and RNA. First, we chose the *NORAD* repeat unit 7 (*NRU7*) as a model template to study the effects of protein binding. This region lies within the well-folded and stable (lowSS) domains identified in the chemical probing experiments (both *in cell* and *ex cellulo*) (**Supplementary Figure 2 and 5**). We further envisioned that choosing such a stable domain would aid in future 3-D structure-determination attempts. Moreover, choosing a larger *NRU78* domain (> 550 nucleotides) would be technically challenging during *in vitro* gel shift experiments. Next, we chose to study the binding of known interactor Pumilio1 and an uncharacterized interactor FUBP3. FUBP3 was chosen not only because it was the most enriched protein in *NRU78* pull-downs in comparison with *MS2* alone controls (**Figure 3C**) but also because previous CLIP experiments have identified overlapping binding sites on *NORAD NRU7* for FUBP3 and PUM1(Munschauer, Nguyen et al. 2018). It is unknown whether FUBP3 competes with Pumilio1 for binding to *NORAD*, or if it cooperatively binds *NORAD* along with Pumilio1. Therefore, FUBP3 serves as an ideal candidate to study the regulatory mechanism of the *NORAD*-Pumilio interaction and thereby its regulation of *NORAD*-associated genome stability maintenance.

We cloned the RNA binding homology domain (HD domain) of human Pumilio1 into a bacterial expression plasmid, recombinantly expressed and purified the protein to high purity and homogeneity (Wang, Zamore et al. 2001) (**Supplemental Figure 10 A and B**). Similarly, we recombinantly expressed and purified the RNA binding K Homology domains (KH domains) 1-4 of human FUBP3 to high purity and homogeneity (Ni, Knapp et al. 2020) (**Supplemental Figure 10 C and D**). Electrophoretic mobility gel shift assay (EMSA) was carried out with purified *NRU7* and PUM1-HD (**Figure 4A**). Multiple copies of Pumilio1 can bind to the *NRU7* as expected. Instead of a stepwise addition, the RNA is bound directly by multiple copies. We do not observe any 1:1 RNP complex formation even at low concentration of Pumilio1 (250 nM). Next, when EMSAs were performed with purified FUBP3 KH1-4 and *NRU7*, a 1:1 RNP complex forms at low protein concentrations (250 nM) (**Figure 4B**). Next, upon increasing the protein concentration, there is a multimeric assembly of FUBP3 on *NORAD NRU7* (corresponding to ∼ 242 kDa to 480 kDa). Finally, EMSAs were carried out where the RNA was first incubated with Pumilio1, followed by addition of increasing concentration of FUBP3 (**Figure 4C**). Addition of Pumilio1 to the naked RNA results in the direct formation of *NRU7*:multiple Pumilio 1 complex formation as observed previously (Lane 2, **Figure 4C**). When FUBP3 is added, there is a super-shift in the *NRU7*-Pumilio1 complex. This shift starts even at addition of 0.12 µM FUBP3 (Lane 3, **Figure 4C**), and this ‘*NRU7*-Pumilio1-FUBP3’ complex is completely formed at 0.37 µM FUBP3 addition (Lane 5, **Figure 4C**). Upon addition of higher concentration of FUBP3, multiple copies of FUBP3 starts to assemble on the *NRU7*-Pumilio1 complex (Lane 6 and 7, **Figure 4C**). Comparison of lanes 2 and 8 with lane 6 shows that, the *NRU7*-Pumilio1-FUBP3 sample runs higher than either the *NRU7*-Pumilio1 (lane 2) or *NRU7*-FUBP3 (lane 8) complex. This suggests that *NORAD* repeat unit 7 (*NRU7)* acts as a platform for assembly of multiple protein subunits such as Pumilio1 and FUBP3.

**Figure 4:**
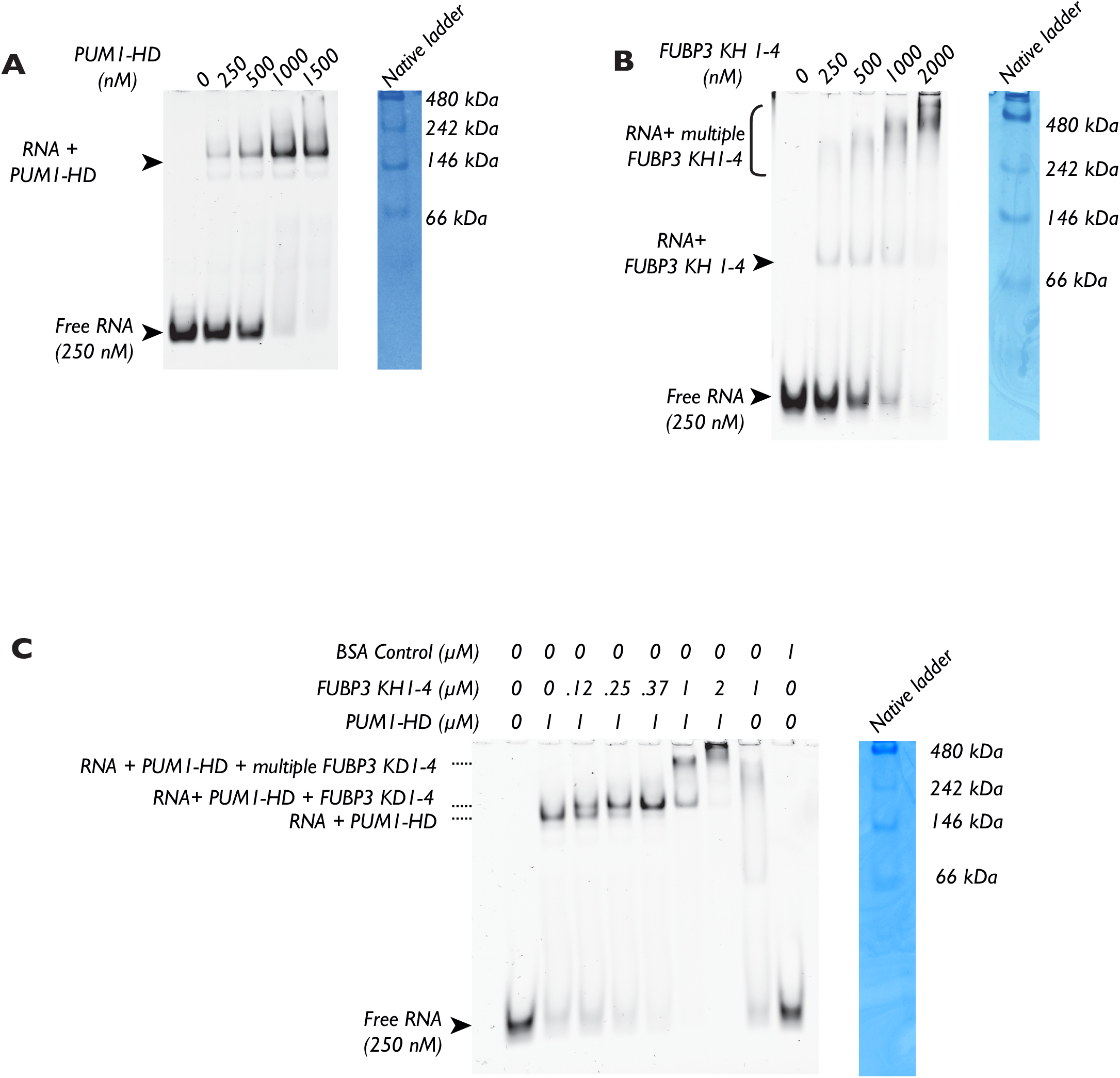
Reconstitution of a minimal *NORAD*-Pumilio1-FUBP3 complex using purified components. **A:** Electrophoretic mobility shift assays (EMSAs) performed with *NORAD* repeat unit 7 (*NRU 7*) RNA with purified human Pumilio 1 HD domain (PUM1-HD). *NRU 7* is 122 nt long and weighs 31 kDa in size. Purified PUM1-HD is 41 kDa. Multiple copies of PUM1-HD appear to bind *NRU7*. **B:** EMSAs performed with *NORAD* repeat unit 7 (*NRU 7*) RNA with purified human FUBP3 KH domains 1 −4. Purified FUBP3 KH 1-4 is 41.7 kDa. It appears that first, *NRU7* has one FUBP3 KH1-4 binding to, and upon increasing the protein concentrations, multiple copies of FUBP3 KH1-4 bind the RNA. **C:** EMSAs performed with *NORAD* repeat unit 7 (*NRU 7*) RNA, purified PUM1-HD and FUBP3 KH1-4. First, the RNA is incubated with 1 µM PUM1-HD, and increasing amounts of FUBP3 KH 1-4 are added to the *NRU 7* - PUM1-HD complex. As a negative control, *NRU7* is incubated with 1 µM BSA. NativeMark™ Unstained Protein Standard (Invitrogen) is used as a ladder to estimate the size of the RNA-protein complexes formed. The samples are run on a 8% native polyacrylamide gel, and RNA bands are visualized by SYBR™ Gold Nucleic Acid Gel Stain (ThermoFisher), and protein bands are visualized by Coomassie staining (AbCam). All EMSAs are done in duplicates. The *NRU7* construct used for EMSA lies within a well-folded (low-Shannon and low-SHAPE) region 2621-2743 nt of *NORAD*.

## DISCUSSION

Using *in cell* SHAPE-MaP pipeline we have successfully probed the secondary structure profile of full-length *NORAD* and discovered up to ten well-folded domains within the RNA. Several of these regions that contain low SHAPE and low Shannon entropy values also overlap with binding sites on *NORAD* for proteins such as RBMX, SAM68 or Pumilio (**Supplemental Figure 3**). Previous studies have shown that the 5′-domain of *NORAD* (1 - 898 nt) lies within the region implicated in binding RNA processing factors such as RBMX and ALYREF (Munschauer, Nguyen et al. 2018). We find that several long stretches (including 14 - 190 nt) within the RBMX binding site appear well-folded and are likely to be protein interaction sites (**Supplemental Figure 2**). Similarly, a recent study showed that regions in the 5′-domain of *NORAD* (1–573 nt) that are rich in G/C content, were also highly structured and remained robust under stress conditions (Ziv, Farberov et al. 2021). Although it has been shown that the 5′-domain of *NORAD* is not required for *NORAD*’s ability to maintain genome stability maintenance, studies suggest that the 5′-region could play a role in *NORAD*’s ability to repress Pumilio mediated mRNA decay by aiding it’s export to the cytoplasm (Elguindy, Kopp et al. 2019, Ziv, Farberov et al. 2021). Future studies will determine the exact mechanism of function of the 5′-domain and the associated biological pathways.

Applying the nextPARS technique to *in vitro* synthesized truncations of *NORAD*, the secondary structure of every *NORAD* repeat unit has been experimentally characterized (Chorostecki, Saus et al. 2021). The study further showed that the binding sites for the SAM68 protein in *NORAD* (that usually lies within *NORAD* repeat units) are robust to structural perturbation. In our *in cell* SHAPE data, we find that SAM68 binding sites in *NRU2* lie within the well-folded domains and further supports the earlier finding that SAM68 binding sites stabilizes the RNA. Additionally, we also find that portions of *NRU6* and the entire *NRU7* fall under our newly discovered well-folded domains (**Supplemental Figure 8**).

A particular region of interest in our study was the unusually long ∼ 200 nt stem region found between the *NRU7* and *NRU8* (78 stem). This region was also recently highlighted in a study from the Ulitsky lab (Ziv, Farberov et al. 2021). Using the COMRADES technique, their study thoroughly characterized the structure of *NORAD*, and discovered several long-range interactions. Particularly, they showed that the structure within the base of this 78 stem is necessary for efficient *NORAD* dependent Pumilio sequestering, possibly by bringing the PREs into close spatial proximity. Next, the regions (3103-3132 nt) immediately after *NRU8* were also identified as well-folded in our study. It is interesting to note that, like the 78 stem, this region also falls immediately after a repeat unit.

Our study surprisingly found that the 3′-domain of *NORAD* (regions after *NRU10*) contains regions of high-SHAPE reactivity (4200-4230 nt, 4240-4310 nt, 4547-5030 nt etc) and these include *NRU11* and *NRU12* (**Supplemental Figure 2 and 3**). Although the nextPARS structure of *NORAD* found that *NRU11* and *NRU12* forms a structured domain, our *in cell* SHAPE data is consistent with previous *in silico* findings (Tichon et al, 2016) and suggests that *NRU11* and *NRU12* do not form the ‘short stem’ and ‘long stem’ that are characteristic of *NORAD* repeat units. Given the high SHAPE reactivity of these regions, it is likely that they are structurally flexible, but they might still form an ordered structure depending on cellular conditions. Using a luciferase-reporter assay in an over-expression setup, it was shown that the 3′-domain of *NORAD* negatively regulates *NORAD* mediated Pumilio derepression (Ziv, Farberov et al. 2021). Given that the 3′-domain of *NORAD* has high SHAPE reactivity, it is tempting to speculate that this region could act as a protein interaction interface, facilitating the recruitment of proteins that negatively regulate *NORAD*-Pumilio interaction. Mechanisms involving the *NRU*s and poly(A) tail cannot be ruled out as well. Additionally, this region could also be involved in pathways independent of Pumilio and needs to be explored.

Along with the two recent structural studies of *NORAD* using nextPARS and COMRADES, our current report probing the structure of *NORAD* in HCT116 cells using a complementary SHAPE-MaP approach serves as a starting point to understand the role of RNA structures within *NORAD*, and will aid in future structure-function studies (Chorostecki, Saus et al. 2021, Ziv, Farberov et al. 2021).

We next utilized *ex cellulo* SHAPE-MaP to discover protein binding hotspots within *NORAD* in an unbiased manner similar to the earlier work on lncRNA *Xist* (Smola, Christy et al. 2016). By comparing the *in cell* and *ex cellulo* reactivities, we identified five protein binding hubs (namely P1 - P5) that were consistently found in all the replicates and across samples (**Figure 2** and **Supplemental Figure 6**). P1 includes the *NRU1* that harbors SAM68 binding sites but no PREs. Regions P2 and P3 lie within the previously identified minimal functional core of *NORAD* (ND4) (**Figure 2)** (Elgundi et al, 2019). It is fascinating to observe that P2 consists of the ∼200 nt long 78 stem that lies in between *NRU7* and *NRU8*, and whose base-pairing interactions were shown to be functionally indispensable for *NORAD*’s function (Ziv, Farberov et al. 2021). However, the loop region at the top part of this 78 stem was previously shown to be dispensable for *NORAD*’s ability to sequester Pumilio. This loop region can be seen in our study to become highly reactive in cell extracts depleted of proteins, suggesting that it could be interacting with non-Pumilio proteins and be involved in other novel pathways (**Figure 2C**). A recent study described the identification of a ‘mini*NORAD*’ that consists of the 5′-domain of *NORAD* plus *NRU78*, and that this construct was sufficient for *NORAD* mediated Pumilio sequestering in cells (Ziv, Farberov et al. 2021). The structural findings presented here showing the discovery of a well-folded protein binding hub within *NRU78*, complements the functional model of mini*NORAD* and further highlights the importance of *NRU78*.

Next, the region P3 that lies immediately after *NRU8* and previously identified as a well-folded region from the *in cell* SHAPE-MaP data, is shown to be a potential protein interaction hub. The stem structures within P3 regions (3090 - 3152 nt) disappear in the *ex cellulo* samples depleted of proteins (**Figure 2D**). Protein interaction hubs P4 and P5 lies after *NRU10* in a well-folded domain region with significantly low SHAPE reactivity in the *in cell* samples. It remains unknown what proteins or cellular factors these hubs might harbor, since they are not known to be involved in Pumilio or SAM68 binding. It is important to note that NAI modifies the 2′-OH of the ribose sugar in flexible regions, and therefore this method cannot identify the footprint of all the proteins that binds structured regions in *NORAD* or of proteins that specifically binds the base of RNA. Therefore, we do not rule out the existence of other protein hotspots within *NORAD*.

The current working model for *NORAD*s function is that the core functional domain of *NORAD* (ND4 or mini*NORAD* containing *NRU7*, the long 78 stem and *NRU8)*, binds multiple Pumilio proteins, nucleates the phase separation and thereby sequesters Pumilio into *NORAD* condensates (Lee, Kopp et al. 2016, Elguindy and Mendell 2021, Ziv, Farberov et al. 2021). The structures in the *NRU78* region, especially the 78 stem, brings together various PREs into close spatial proximity, therefore likely enabling *NORAD*-Pumilio condensation. We hypothesize that in addition to RNA structure within this core *NRU78* region, binding of accessory protein partners to regions P2 or P3 could regulate the *NORAD*-Pumilio interaction, and thereby control cellular fate by modulating Pumilio sequestration or release. Therefore, it was essential to identify these protein partners to fully understand *NORAD*-Pumilio dependent genomic maintenance.

Using an *in vitro MS2* based pull-down approach, we isolated *NORAD NRU78* associated RNPs from HCT116 cell extracts (**Figure 3A and 3C**). Although previous studies have characterized the protein partners of *NORAD* domains (Munschauer, Nguyen et al. 2018, Spiniello, Knoener et al. 2018, Tichon, Perry et al. 2018), here we used the native purification pipeline to ensure that the RNA is maintained under non-denaturing conditions throughout the entire experiment (**Supplemental Figure 8A**). Interestingly, amongst the highly enriched proteins in the *NRU78* samples were FUBP3, HARS2, CELF1 and PUM1 (**Figure 3C and Supplemental Figure 9C**). In order to precisely identify the factors that bind the loop in the 78 stem region (**Figure 2C**), we designed a *NRU78* RNA with a mutated loop region as an additional control in the pull-downs (**Figure 3B**). Comparing the protein enrichment levels between the wild-type *NRU78* and *NRU78* loop mutant, led us to discover several mitochondrial ribosomal proteins as potential interactors of this region (**Supplemental Figure 9D**). HARS2, a mitochondrial aminoacyl-tRNA synthetases was also found enriched in these comparisons suggesting that the binding site of HARS2 in *NRU78* is unambiguously the tip of the 78 stem loop. Given the fact that several mitochondrial ribosome associated proteins are enriched in this region and that recent studies have shown that this loop is dispensable for *NORAD*-Pumilio interaction, it is tempting to speculate that the 78 stem loop in *NORAD* could be involved in additional pathways independent of Pumilio function, possibly related to protein synthesis. Further studies are necessary to clarify this.

While *NORAD*’s association with Pumilio has been extensively studied, the roles of these newly discovered factors with regards to *NORAD* remain uncharacterized. Previous CLIP studies have shown that FUBP3 and FUBP1 occupy binding sites on *NORAD* throughout its sequence (except the 5′-domain), and their binding sites show a strong peak on *NORAD* near NRU7 (Munschauer, Nguyen et al. 2018). This peak also coincides with PUM1 CLIP peaks (Munschauer et al, 2018). The FUBP family of KH domain proteins have been extensively characterized for their involvement in activating the transcription of the *MYC* oncogene (Avigan, Strober et al. 1990, Duncan, Bazar et al. 1994). It has also been shown that even minute changes in the expression of *MYC* oncogene can alter cellular fate (Shichiri, Hanson et al. 1993). Could it be possible that *NORAD* sequesters FUBP3 proteins in the cytoplasm to prevent their transport into the nucleus, similar to how *NORAD* sponges Pumilio proteins? It has been shown that *NORAD* forms phase-separated condensates with Pumilio in the cytoplasm (Elguindy and Mendell 2021). It is well known that such membrane less condensates contain multiple proteins, the composition of which changes based on the cellular environment (Mitrea and Kriwacki 2016). Moreover, proteins such as Pumilio that enter these condensates often do so via a large unstructured and disordered domain (Borcherds, Bremer et al. 2021, Elguindy and Mendell 2021). Domain predictions of FUBP3 suggest that the C-terminal ends beyond amino acid residue 334 are largely disordered. It is therefore possible that besides Pumilio, *NORAD* condensates in the cytoplasm can act as a reservoir for sequestering other RNA binding proteins containing disordered domains such as FUBP3 or CELF1, therefore regulating multiple cellular pathways. Alternatively, some of these newly discovered factors could compete with Pumilio to bind *NORAD* at *NRU78*, and subsequently release Pumilio from the phase separated condensates depending on the cellular state.

To test the above hypotheses, we adopted an *in vitro* reconstitution system with pure recombinant proteins and purified *NORAD* domains. As a model RNA, we chose *NORAD NRU7* due to its low Shannon-low SHAPE property and its proximity to the functionally relevant core of *NORAD*. Next, we chose to focus on FUBP3 due to its overlapping binding sites on *NORAD* with PUM1. Our EMSA experiments shows that *NRU7* can indeed bind multiple copies of PUM1, and multiple copies of FUBP3 when each is added to the RNA separately. Interestingly, the formation of *NRU7* - multiple PUM1 complexes even at lower concentration of PUM1 suggests that multiple PUM1s can simultaneously assemble on naked RNA (**Figure 4A**). Upon increasing the concentration of PUM1, the molecular size of the RNP remains the same, suggesting that the RNA binding domains of PUM1 do not enable the formation of higher order oligomers (**Figure 4A**). Next, when we do the EMSAs with FUBP3, we first see the appearance of a *NRU7*-FUBP3 1:1 complex (at lower concentration of FUBP3), and at higher concentrations of FUBP3, we see multimeric FUBP3 assembly on *NRU7*. We know that *NRU7* is specifically recruiting PUM1 and FUBP3 since the control lanes with BSA remain unchanged. Our EMSAs with both PUM1 and FUBP3 added to the RNA suggests that *NRU7* first forms a stable complex with PUM1 that in turn acts as a platform for the assembly of higher order oligomers of FUBP3. Future studies aimed at characterizing the functional significance of *NRU7*-Pumilio1-FUBP3 complex assembly are essential.

In summary, using an integrative structural biology approach that combines structure probing, and proteomics, we discovered a well-folded protein binding hub within the core of *NORAD* i.e. *NRU78* and identify the protein partners of this region in an unbiased manner. *In vitro* biochemical reconstitutions suggest the formation of a higher-order protein-RNA assembly in this region and further studies are needed to clarify the exact roles these factors play in the functioning of *NORAD*. We hope that the structural and proteomics data presented in this study will be a valuable resource for future mechanistic studies of *NORAD*.

## LIMITATIONS

This report has been prepared with the aim of making available to the public, the structural and proteomics data generated in this study. Future experiments are necessary to validate the functional relevance of these structural findings inside the cells, and to understand the roles of these newly discovered protein partners in *NORAD*s function. EMSAs were performed with purified RNA binding domains of relevant proteins, and not the full-length constructs.

## AUTHOR CONTRIBUTIONS

A.K., A.M.P. and H.W. designed the experiments. A.K., H.W., S.P. and R.T. performed and analyzed the SHAPE experiments. A.K. performed the RNA-protein pull downs. Mass spectrometry sample preparation and data collection were carried out in collaboration with Creative Proteomics Pvt. Ltd. (New York). H.W. analyzed the mass spectrometry data. A.K. and Z.P. purified proteins. A.K., Z.P., and S.P. performed EMSAs. This report was written by A.K. with inputs from A.M.P. and H.W.

## ACKNOWLEDGEMENT

## Acknowledgements

The work was supported by HHMI awarded to A.M.P., the Human Frontier Science Program Long-Term Fellowship (HFSP) awarded to A.K. and the China Scholarship Council (CSC)-Yale World Scholars Program in Biomedical Sciences awarded to H.W.. Z.R.P. is supported by the National Science Foundation Graduate Research Fellowship Program (DGE-2139841). The authors also acknowledge Dr. Mathias Munschauer at the Helmholtz Institute for RNA-based Infection Research for providing pDONR221-*NORAD* plasmid.

## MATERIALS AND METHODS

### RNA structure probing

HCT116 cells (passage #7) were seeded into McCoy’s 5A Medium (ThermoFisher) supplemented with 10% heat-inactivated FBS and 1x Antibiotic-Antimycotic (ThermoFisher) in 150 mm diameter sterile tissue culture plates, and grown at 37°C. 48 hours post seeding, the growth medium was removed and the cells were washed with sterile PBS. The adherent cells were dislodged using a cell-scraper, and resuspended in 4 mL PBS with a final cell count of 4.0 × 10^6^ cells/mL. For the *in cell* SHAPE reaction, to 0.9 mL of this cell-suspension, 100 µL of 2M NAI (MilliporeSigma)(final concentration of 200mM) was added, mixed well and incubated at room temperature for 10 mins. In parallel, as a control, 100 µL of DMSO was added to another tube containing 0.9 mL of the cell re-suspension. 3 mL Trizol (ThermoFisher) was added to 1 mL of cellular PBS suspension in a 15 mL test tube, and incubated at room temperature for 2 mins. Total RNA extraction was carried out by the addition of 1 mL of chloroform:isoamyl alcohol (24:1). The aqueous phase was separated by centrifugation at 15,000g for 15 mins. RNA was precipitated from the aqueous phase by the addition of 8 mL of cold 100% ethanol, mixing the suspension and leaving it in the - 80°C for 16 hours. Upon centrifugation at 12,000g for 20 minutes, the RNA pellet becomes visible. The pellet is washed once with 70% ethanol, before being dissolved in 490ul 1x TURBO™ DNase buffer (ThermoFisher). Add 5 µL of SUPERaseIn™ RNase Inhibitor (Invitrogen) at 20 U/μL to the resuspended RNA. Next, 5 µL of TURBO™ DNase (ThermoFisher) was added to the RNA and was incubated at 37°C for 15 minutes. Another 5 µL of TURBO™ DNase (ThermoFisher) was added to the tube, and incubated for 15 minutes. The RNA was further cleaned by using an RNeasy MinElute Cleanup Kit (Qiagen), and eluted in 20 uL of ME buffer (8mM MOPS, 0.1mM EDTA, pH 6.5) supplemented with 1 µl of SUPERaseIn™ RNase Inhibitor (Invitrogen) and used in reverse transcription (RT) reactions.

In case of *ex cellulo* SHAPE probing, HCT116 cells were grown as described above, and resuspended in 4 mL PBS. Next, the cells were gently centrifuged at 1000 g for 5 mins at 4°C and the PBS (supernatant) was removed without disturbing the cell pellets. The cells were next, resuspended in 4mL of cell lysis buffer (50 mM HEPES-KOH pH 7.5, 150 mM KCl, 6 mM MgCl2, 1mM CaCl2, 10 mM Ribonucleoside Vanadyl Complex (NEB), 0.2% v/v Triton X-100and 0.12 U/μL SUPERase•In™ RNase Inhibitor). To 1 mL of this suspension, 20 µl of TURBO™ DNase (ThermoFisher) was added and incubated for 15 minutes at 37°C. Next, 40 µl of Proteinase K (ThermoFisher) was added to this tube and incubated for 15 minutes at 37°C. The tube was then clarified by centrifugation at 600g for 30 seconds, and the ∼ 0.9 mL of the supernatant was transferred to a fresh tube. SHAPE probing was done by adding 100 µl of NAI and the subsequent steps were similar to above described *in cell* SHAPE protocol.

### Reverse Transcription and amplicon <u>generation</u>

Eleven ∼ 600 nt amplicons tiled across the entire sequence of *NORAD* were designed to obtain full sequencing coverage. Each of these amplicons had an overlapping ∼ 200 nucleotides with their adjacent counterparts in order to ensure we obtained data for the primer binding sites. All the primers used in RT were designed using OligoWalk (Lu and Mathews, 2008). This helped avoiding highly-structured primers and binding to highly-structured regions in *NORAD*. The RT primers used in the study are listed in **Supplementary Table 2**. While designing the amplicon PCR primers, the reverse primers were inset 3 nt from the 5′-end of the corresponding RT primer. For each RT reaction, 1500 ng of total RNA was used as input. 2 µl of 1 µM RT primers were added to 8 µl of total RNA. Annealing reaction was done at 72°C for 2 minutes and then cooled to room temperature, followed by addition of 8μL of 2.5x SuperScript™ II Reverse Transcriptase buffer (ThermoFisher), 1.5 µl of milliQ water and 0.5 µl of SuperScript™ II Reverse Transcriptase. The RT reaction is done at 42°C for 3 hours. The generated cDNA was cleaned using ampure xp beads (beckman coulter), and eluted in 10 µl milliQ water (nuclease free). Amplicons were generated using Q5® High-Fidelity DNA Polymerase (NEB) using *NORAD* specific forward and reverse primers, and 5 µl of cDNA per reaction carried out in a 50 µl total volume per amplicon.

### Library preparation and sequencing

The amplicons from the Q5 PCR step were cleaned up using Monarch® PCR & DNA Cleanup Kit (NEB). The purity and size of the DNA bands were visualized by running a 1% agarose gel. The amplicons were next diluted to a concentration of 0.2 ng/ µ. The diluted amplicons were separated into either odd or even pools. The NexteraXT DNA Library Preparation Kit (Illumina) was used to generate sequencing libraries (1/5th of the recommended volume). Qubit dsDNA HS Assay Kit (ThermoFisher) and BioAnalyzer High Sensitivity DNA Analysis (Agilent) were used to determine library concentration and their average size. Libraries were then diluted to 4nM, denatured, and the final library was generated following Illumina’s protocols. Sequencing was done using a NextSeq 500/550 platform and a 150 cycle mid-output kit.

### SHAPE-MaP data processing and structure predictions

ShapeMapper 2 (Busan and Weeks, 2018) was used to process all the obtained libraries in this study. The reads were aligned to the full length human NORAD sequence (Gene ID: 647979). A minimum of 5000x read-depth was used as a quality control benchmark. Mutational rates were calculated between NAI-treated and DMSO-treated samples similar to previous studies (Guo, Adams et al, 2020, JMB). The SHAPE reactivities from SHAPEmapper2 were used as constraints to generate predicted structures for the entire *NORAD* RNA using SuperFold (Smola et al., 2015). The Shannon entropy values were calculated as described previously (Houston, Wan et al, 2021). The output structures (.ct files) from SuperFold were visualized in StructureEditor, in RNAStructure package (Reuter and Mathews, 2010). The corresponding SHAPE reactivity values were superimposed on the predicted structure to draw the secondary structures reported in this study. To discover well-folded regions in the RNA, the low-Shannon and low-SHAPE reactivity metric was used as described previously (Houston, Wan et al, 2021; Weeks, 2021; Smola et al, 2016). For each condition, two replicate datasets were used. Local median SHAPE reactivity and Shannon Entropy values were determined in 50nt sliding windows. The global median SHAPE reactivity or Shannon Entropy were then subtracted from calculated values to better visualize the well-folded regions.

### Protein Expression and Purification

#### MBP-MS2

pMBP-MS2 (Plasmid #65104, Addgene) was transformed into BLE1 DE3 cells. 2 L of LB (with 50 ug/mL Amp) with 10 ml of the overnight culture were grown at 37°C until ∼0.5 and 4mL of 100mM IPTG was added to each 1L of the culture. The cell growth was continued at 21°C for 18 hours at 180 rpm, following which the cells were harvested by centrifugation. The cell pellet was resuspended in 50 mL of ice cold lysis buffer (20 mM HEPES, pH 7.9, 200 mM KCl, 1 mM EDTA, one tablet of Protease inhibitor tablet / 100 mL). Once the cells are completely re-suspended, transfer the suspension into a 100 mL glass beaker, and lyse the cells using ultrasonication (BIG tip, 35% amplitude, total of 3 minutes with 30 seconds ON and 60 seconds OFF). The lysed cells are clarified by centrifugation in 50 mL falcon tubes at 30000 g for 40 minutes. The clarified lysate is mixed with 1 mL of amylose resin (E8021S, NEB) pre-equilibrated with lysis buffer and incubated in the cold room using an end-over-end rotor for 2 hours. After incubation, the unbound proteins / flow through are removed by spinning down the resins at 600g. The resins are then washed with 80 ml of 20 mM HEPES, pH 7.9, 20 mM KCl, 1 mM EDTA. Next, 1 mL of elution buffer (20 mM HEPES, pH 7.9, 20 mM KCl, 1 mM EDTA, 10 mM maltose) is added to the washed resins, mixed well with a pipette, and the elution fractions collected immediately. In total, twenty 1mL elution fractions were collected. The elution fractions were pooled, and concentrated using a 30kDa amicon centrifugal filter up to a volume of ∼ 1.2 mL. At this stage, the sample might contain nucleic acid contamination. To clean the MBP-MS2 protein further, we use a 1 mL HiTrap® Heparin High Performance column (GE17-0407-01). After loading the sample onto the Heparin column, the column is washed with 10 column volumes of heparin buffer A (20 mM HEPES, pH 7.9, 20 mM KCl, 1 mM EDTA). The MBP-MS2 protein is eluted by running a gradient from 20 to 400 mM KCl over 10 column volumes. The elution and wash fractions are run on an SDS-PAGE to assess sample purity and check for presence of any degradation factors. Fractions containing the protein at the right size and purity are concentrated using an amicon 30kDa cut off up to a concentration of 10 mg/mL. The final protein is in a buffer containing 20 mM HEPES, pH 7.9, 20 mM KCl, 1 mM EDTA.

#### PUMILIO1 RNA binding domain (PUM1-HD)

The regions encoding the RNA binding HD domain of human Pumilio 1 gene (Gly-828 to Gly-1176) were cloned into pGEX-6P-1 using the following primers:

Fwd_BamHI_PUM1_HD: 5′-*CGCGGATCCATGGGCAGGAGCAGGCTTTT*-3′ & Rev_ECORI_PUM1_HD: 5′-C*CGGAATTCTTATTACCCTAAGTCAACACCGTTCT* −3′

Standard restriction enzyme based cloning using the BamHI and EcoRI sites were used to perform the cloning. The resulting protein expression plasmid contained the PUM1 HD gene with a N-terminal GST-tag followed by a 3C PreScission protease site. Protein expression and purification were done similar to previously described methods (Webster et al, 2019). Briefly, 4L of BL21 DE3 cells (containing pGEX-6P-1_PUM1-HD) were lysed by sonication in ∼140 mL lysis buffer (50 mM Tris-HCl pH 8, 250 mM KCl, 1 mM TCEP) supplemented with protease inhibitor cocktail (Roche) and DNase I (5 μg/ml) (Sigma). Lysis was done using ultrasonication (BIT tip, 30 sec ON, 60 sec OFF) for 3 mins in total. The lysate was cleared by centrifugation at 30,000 × g for 40 min at 4°C. Clarified lysate was applied to Glutathione Sepharose 4B affinity resin (1 ml bed volume per 2 l culture; New England Biolabs), and incubated with rotation for

1.5 hr at 4°C. Resin was separated from lysate by spinning down at 500g for 5 mins and washed two times with 50 ml of GST wash buffer (20 mM Tris-HCl pH 8, 50 mM KCl, 1 mM DTT). Protein was eluted from the resin in 5 ml of glutathione elution buffer (75 mM HEPES pH 7.4, 300 mM NaCl, 5 mM DTT, 50 mM glutathione) by incubation at room temperature and rotation for 10 minutes per elution fractions. The elution fractions were pooled together in a 50 mL falcon tube and concentrated into a 4 mL volume. The elution was next topped upto 20 mL with cleavage buffer (20 mM Bis-Tris pH 6.5, 50 mM NaCl, 1 mM DTT, 10% Glycerol) and 400 uL of home-made 3C protease was added to the tube. The GST-cleaved PUM1-HD sample was next loaded onto an ion exchange chromatography using a 5 mL Heparin column (GE) equilibrated at a salt concentration of 100mM. Pure PUM1-HD was eluted by doing a shallow linear gradient upto 1M NaCl across 8 column volume. This step ensured the removal of nucleic acid contaminants. The purified protein was then polished by a size exclusion chromatography step using a Superdex ® 200 Increase 10/300 GL (Sigma) in storage buffer (10 mM Tris-HCl pH 7.4, 150 mM NaCl, 5 mM DTT). The peak fractions were then run on an SDS-PAGE to check the purity and assess the size. The pure peak fraction from SEC were pooled together, concentrated to 70 µM, flash frozen in liquid nitrogen and stored at −80 °C.

#### FUBP3 RNA binding domains (FUBP3 KH1-4)

The KH domains 1, 2, 3 & 4 of human FUBP3 (residues 75-427) were cloned into a pET-28 a (+) vector using the following primers:

FP_FUBP3_KH1-4_BamHI: 5′-*GATGATCGGATCCATGaggacggtaataacggaa* - 3′ RP_FUBP3_KH1-4_HindIII: 5′-*CCCAAAGCTTTTATTAcccgccaactttctcatcta* - 3′

Standard restriction enzyme based cloning using the BamHI and EcoRI sites were used to perform the cloning. The resultant pET-28 a (+) _FUBP3_KH1-4 plasmid contained a N-terminal 6-His tag followed by the FUBP3_KH1-4 gene sequence. The expression plasmid was transformed into BL21-CodonPlus Competent Cells (Agilent), and protein expression was done by the addition of IPTG (1 mM final concentration) and growing the cells at 16 °C for 14 hours. Protein purification was done similar to previously described methods (Ni et al, 2020). Briefly, pellets from 2 L of cells were resuspended in 160 mL of cold lysis buffer (50 mM HEPES pH 7.5, 500 mM KCl, 20 mM Imidazole, 1 mM TCEP, 5% glycerol, 4 µg/µl DNAseI, protease inhibitor). The resuspension is then sonicated using an ultrasonicator (BIG tip, 40% Amp, 10 sec ON, 15 sec OFF, 20 mins ON in total). Sonication is done until the opacity of the resuspension goes from being turbid to somewhat clear. The lysate was then clarified by centrifugation at 15000 g for 30 mins. 2mL of Ni-NTA resins (Qiagen) equilibrated with lysis buffer, were mixed with the supernatant from the centrifugation step i.e., the clarified lysate. The resin-lysate suspension is incubated in the cold room rotor for 1 hour. The mixture is spun down at 600g for 5 mins, the supernatant containing the unbound proteins are removed. The resin is washed with 100 mL of wash buffer (50 mM HEPES pH 7.5, 300 mM KCl, 20 mM Imidazole, 1 mM TCEP). The proteins are eluted by adding 5 mL of cold elution buffer (50 mM HEPES pH 7.5, 300 mM KCl, 300 mM Imidazole), and collecting the flow through using a gravity column. Ten elution fractions are collected. The eluted fractions are pooled together, concentrated, and diluted to 50 mL using HEPES pH 7.5 so that the final concentration of salt in the sample is 50 mM KCl. The samples are next loaded onto a 5 mL Hitrap S column (Qiagen) pre-equilibrated with 50 mM HEPES pH 7.5, 50 mM KCl, 0.5 mM TCEP. An external 50 mL superloop was used to load the sample. The flow was done at 0.5 mL/min and the elution was carried out by a linear gradient of KCl upto 1M concentration over 5 column volumes. 0.5 mL elution fractions were collected. The peak fractions from the S-column were concentrated, and loaded onto a Superdex 200 10/300 Increase GL column (Sigma) using a 500 uL sample loop and run at 0.5 mL / min. SEC buffer was 20 mM HEPES pH 7.5, 150 mM KCl, 0.5 mM TCEP. The peak fractions were then run on an SDS-PAGE to check the purity and assess the size. The fraction containing the majority of pure proteins were pooled and concentrated to 259 µM, flash frozen in liquid nitrogen and stored at −80 °C.

### Cloning

RNA constructs of interest were amplified from a parent plasmid (gene blocks from ThermoFisher) and inserted into a linearized pcDNA3.0 using Gibson assembly. We introduced the RNA coding sequence in between The T7 start site and the EcoRI site in pcDNA3.0.

The following primers were used to linearize pcDNA3.0:

pcDNA3 FP1: 5′-*gaattctgcagatatccatcacactgg* −3′

pcDNA3 RP1: 5′-*CCCTATAGTGAGTCGTATTAATTTCGATAAGC* −3′

A gene block containing *NORAD NRU 7&8* sequences from 2621 nt – 3067 nt, immediately followed by a triple MS2 stem-linker sequence (PNAS September 17, 2002 99 (19) 12203-12207) was purchased from ThermoFisher. This was used as a template to make *pcDNA3.0_NORAD_NRU78_MS2.* To make the *pcDNA3.0-NORAD_NRU78_LoopMutant_MS2, a gene block containing* 5′-*ATGGATACGAATGCAGATGTAAAGAGGCTGATGTCCTGCAGTTTAAGGGGAAATGGAATACCTAGCGTGGCGGAGTCGAATTTTGACGA -* 3′ instead of 5′-*AAGTGAGATAACATCAGCTCTAAGTGACACGTGCCTATATCCATCAGGTTGGTGGTGGAGAGGAGTTGGAAGGAATGAAGGGTTCTAGA-* 3′ within in the 78 Stem loop of *NRU78* was ordered, immediately followed by a triple MS2 stem-linker sequence.

The DNA sequence encoding *NRU 78_MS2* or *NRU78_LoopMutant_MS2* was amplified from the above gene blocs by PCR using following primer sets:

Norad_FP1: 5′-*gcttatcgaaatTAATACGACTCACTATAGGG-* 3′

Norad _MS2_RP2: 5′-*CCAGTGTGATGGATATCTGCAGAATTCCGTACCCTGATGGTG* −3′

The entire sequence of *NORAD NRU 78* was fed into the SMS Shuffle tool to generate a random sequence that has the same base composition and size as that of *NRU 78*. This randomer followed by a triple *MS2* stem was purchased from ThermoFisher as a gene block. To make Gibson inserts for this construct, the following primers were used:

NRU78 mutant or Randomer shuffle FP1: 5′-*gcttatcgaaatTAATACGACTCACTATAGGG*-3′ Randomer Shuffle RP1: 5′-*GATGGTGTACGGATATCGGATCCCGTTAGAATCTATTTGGTAAATACTCAGTTG* −3′

Reverse complement (order) (54 nt 41% GC Tm: 73°C)

A gene block containing three tandem MS2 stem loop sequence (5′-*CGGGATCCGATATCCGTACACCATCAGGGTACGAGCTAGCCCATGGCGTACACCATCAGGGTACGACTAGTAGATCTCGTACACCATCAGGGTACGAATTCTCTAGAGC*-3′) was used as a template to amplify the triple MS2 stem using the following primers:

MS2triple_FP4: *5*′*-CAACTGAGTATTTACCAAATAGATTCTAACGGGATCCGATATC-3*′

MS2triple_RP4: 5′-*ccagtgtgatggatatctgcaGAATTCCGTACCCTGATGGTGTACGAGATCTACTAGT*-3′

The PCR products containing the aforementioned various RNA constructs were cleaned using a PCR DNA clean up kit (Zymo Research), and used in Gibson assembly reactions (5x Gibson buffer: 1M Tris-HCL pH 7.4, 1M MgCl2, 10 mM dNTPs, 1mM DTT, 25% PEG-8000, 50 mM NAD,. The Gibson reaction was done at 50 °C for 1 hour.

### RNA purification

Plasmids *pcDNA3.0_NORAD_NRU78_MS2, pcDNA3.0-NORAD_NRU78_LoopMutant_MS2, pcDNA3.0_Randomer_MS2 and pcDNA3.0_MS2* were linearized by restriction digestion using EcoRI. 15 µg of linearized pcDNA3 RNA plasmids were used as a DNA template in 500 µl *in vitro* transcription (IVT) reactions. IVT was carried on for 1 hour at 37 °C, after which 4 µl of 1M MgCl_2_ was added to it. After 1 more hour of the reaction, 10 µl of RNase-free DNase I (NEB) was added to stop the reaction. Finally, 12 µl of 0.5 mM EDTA was added to the sample, and the sample was buffer exchanged into the Size exclusion chromatography (SEC) buffer (50 mM HEPES pH 7, 150 mM KCl, 0.5mM EDTA) in a fixed-angle rotor at 3000 g for 20 mins using an amicon 50 kDa cut off (Millipore). 500 µl of concentrated RNA sample is loaded onto a Sephacryl 400 SEC column at room temperature. The peak fractions from the chromatography step are pooled together after assessing their purity and confirming the size by running a denaturing Urea-polyacrylamide gel.

To prepare the RNA substrate used in EMSAs, the following primers were used to amplify the well-folded (low-Shannon & low-SHAPE) 122 nt regions in *NORAD* NRU 7 (2621-2743 nt). This construct encompasses almost the entire NRU7 and contains two PREs and will serve as an ideal substrate to study *NORAD*-protein interactions.

Norad_FP1: 5′-*gcttatcgaaatTAATACGACTCACTATAGGG*-3′

NRU7_S1_RP2: 5′-*TTTACTAACTCTACTTCTGTCATACATTG* −3′

The PCR product is then used as a template for *in vitro* transcription reaction similar to the aforementioned protocol. The IVT product is run on a 5% denaturing urea polyacrylamide gel, and the RNA band of appropriate size is excised and purified using gel-extraction techniques (Timothy W. Nilsen, CSHL protocols). Pure NRU7 RNA is eluted in 400 μl ME buffer supplemented with 5 μl of RNAseOUT (ThermoFisher) at a final concentration of 38 μM, and used for further biochemical studies.

### *In Vitro* RNP pull down

3 mL of purified RNA sample was folded at 37 °C in 50 mM HEPES pH 7, 150 mM KCl, 3 mM MgCl_2_. While the folding reaction is taking place, 180 µl of amylose resin is washed with 5 mL of wash buffer I (20 mM HEPES, pH 7.9, 100 mM KCl, 3mM MgCl2, 1 mM TCEP, 0.01% Tween-20, 10 U RNAseOut). Add purified MBP-MS2 protein to the washed resins to a final concentration of 2 µM, and the binding reaction is carried out for an hour in the cold room. Wash the excess MBP-MS2 using wash buffer I. 720 µL of wash buffer (+ 80 uL of 10X protease inhibitor + 20uL of RNAseOUT) are added to the resin. 20 µL of this slurry is taken as input for the SDS-gel. 110 µl of the slurry was transferred into a fresh tube, 3 mL of the folded RNA added to the slurry, and the tube was topped up with 5µl of RNAseOUT. RNA binding was done for 30 mins at room temperature in an end over end rotor. Next, the unbound RNA was removed by gentle centrifugation at 600g for 5 mins. The resins were washed with 2 mL wash buffer II (20 mM HEPES, pH 7.9, 100 mM KCl, 3mM MgCl2, 1 mM TCEP, 0.01% Tween-20, 0.2 % NP-40, 1X protease inhibitor, 10 U RNAseOut). The resin is then re-suspended in 240 uL of wash buffer II. 10 uL of slurry is taken as an input RNA sample for denaturing Urea-polyacrylamide gels. While the MBP-MS2 protein is being incubated with the amylose resin, the cell lysate is prepared. Each pull-down experiment (for each RNA construct) was performed with lysate coming from ∼ 50 × 10^6^ HCT116 cells. 50 × 10^6^ HCT116 cells are resuspended in 2 mL of cold lysis buffer (10 mM Tris-HCl pH 8, 100 mM KCl, 2 mM MgCl_2_, 0.3 M Sucrose,0,2% v/v Igepal (NP40 CA 630), supplemented with protease inhibitors). The cell slurry is incubated on ice for 5 minutes, and the cells are passed through a 30 Gauge 1/2 Inch 13mm sterile disposable needle fifteen times. The syringe and needles are pre-chilled prior to the lysis. The lysate is clarified by centrifugation at 20,000g for 5 minutes. The clarification step was repeated at least three times to ensure that the viscous lipid-like layer and the cell debri-pellet were avoided while separating the supernatant. Finally, the clarified supernatant was passed through a pre-chilled 0.45 µ sterile filter to remove any additional debris that might otherwise occlude protein-RNA complex formation on the amylose resins. Next, 1 mL of total cell lysate was added to the tubes containing RNA-MBP-MS2 loaded amylose resins, and incubated for 90 minutes in the cold room using an end-over-end rotor. The unbound proteins were removed by centrifugation at 600g for 10 minutes. The resins were next washed with 0.5 mL of wash buffer II. RNA-protein complex (RNP) elution was done by adding 50 µl elution buffer (20 mM HEPES, pH 7.9, 100 mM KCl, 3mM MgCl2, 10 mM maltose, 1 mM TCEP, 0.01% Tween-20) and incubating the slurry for 10 minutes on ice, followed by gentle centrifugation at 600g. 10 µl of elution fraction is taken for running the SDS-PAGE and UREA-PAGE gels. The elution step is repeated for a second time to collect 50 µl more of RNPs. The elution samples were flash frozen in liquid nitrogen and stored at - 80 °C until further use.

### Mass spectrometry and data analysis

Mass-spec experiment and analysis were conducted by Creative Proteomic (New york) using the nanoLC-MS/MS platform. Briefly, samples were purified and digested using a 12% separating gel before being loaded onto the Ultimate 3000 nano UHPLC system (ThermoFisher Scientific, USA). 1 μg of samples were loaded (mobile phase: A: 0.1% formic acid in water; B: 0.1% formic acid in 80% acetonitrile). Full scan was performed between 300-1,650 m/z at the resolution 60,000 at 200 m/z, the automatic gain control target for the full scan was set to 3e6. The MS/MS scan was operated in Top 20 mode using the following settings: resolution 15,000 at 200 m/z; automatic gain control target 1e5; maximum injection time 19ms; normalized collision energy at 28%; isolation window of 1.4 Th; charge sate exclusion: unassigned, 1, > 6; dynamic exclusion 30 s. Subsequently, the raw MS files were analyzed and searched against a human database utilizing MaxQuant. Label-free quantitation (LFQ) values from each MaxQuant output were imported into R for further downstream analysis. The R package ‘Differential Enrichment analysis of Proteomics data’ (DEP) was employed to identify enriched proteins in different samples. Data filtering, normalization, and imputation were performed using the default DEP workflow. Enriched proteins were defined using a cutoff of log_2_ fold change > 1.5 and a P-value < 0.05.

### Gel-shift assays

The gel shift assays were carried out in EMSA buffer (20 mM Tris-HCl pH 8, 100 mM KCl, 0.5 mM TCEP, 5 µl of RNAseOUT/5 mL buffer) and in 20 µl volume. Briefly, 4 µl of purified RNA (NRU7) was added to 16 µl of recombinant proteins so that the final concentration of RNA is 250 nM, and the protein concentration were either 250nM, 500 nM, 1 µm or 2 µM. Accordingly, substocks of proteins were made from the parent stock of 70 µM PUM1-HD and 259 µM FUBP3 KH1-4. All dilutions were performed in the EMSA buffer. The binding reaction was carried out on ice for 1 hour. In case of the multiple protein binding experiment, the RNA was incubated with appropriate concentration of PUM1-HD (final concentration in 20 µl volume equals 1 µM), incubated on ice for 45 mins, and additional FUBP3 KH1-4 was added at appropriate concentrations (final concentration in 36 µl volume equals either 0.125 µM, 0.25 µM, 0.375 µM, 1 µM or 2 µM). In the control experiment, EMSA buffer was added to the RNA sample, incubated for 45 minutes, followed by the addition of Bovine Serum Albumin (Sigma) to a final concentration of 1 µM. The second protein addition was followed by further incubation on ice for 45 more minutes. A native loading dye containing 50% glycerol and 0.1 %w/v bromophenol blue was added to the binding reaction. 14 µL of the loading dye-RNP mixture were loaded onto the wells of a 8% native acrylamide gel (1.5 mm thickness). The gel was run at 5W for 2 hours in a 1x TBE buffer, and in a cold room. To visualize the RNA bands, the gel was first stained using SYBR™ Gold Nucleic Acid Gel Stain (ThermoFisher) for 15 minutes, washed with milliQ water for 30 minutes and imaged in an Amersham™ Typhoon™ Biomolecular Imager (Cytvia) using the Cy3 filter. Once the RNA bands are successfully visualized, protein bands are visualized by re-staining the same gels by Coomassie staining (AbCam) overnight. This is followed by incubation with milliQ water for 4 hours, and subsequent imaging in an Amersham™ Typhoon™ Biomolecular Imager (Cytvia) using the Cy5 filter.

## DATA AVAILABILITY

All chemical probing data are available upon request.

