## Supplemental Figure 1 for "Discovery of a Well-Folded Protein Interaction Hub Within the Human Long Non-Coding RNA *NORAD*"

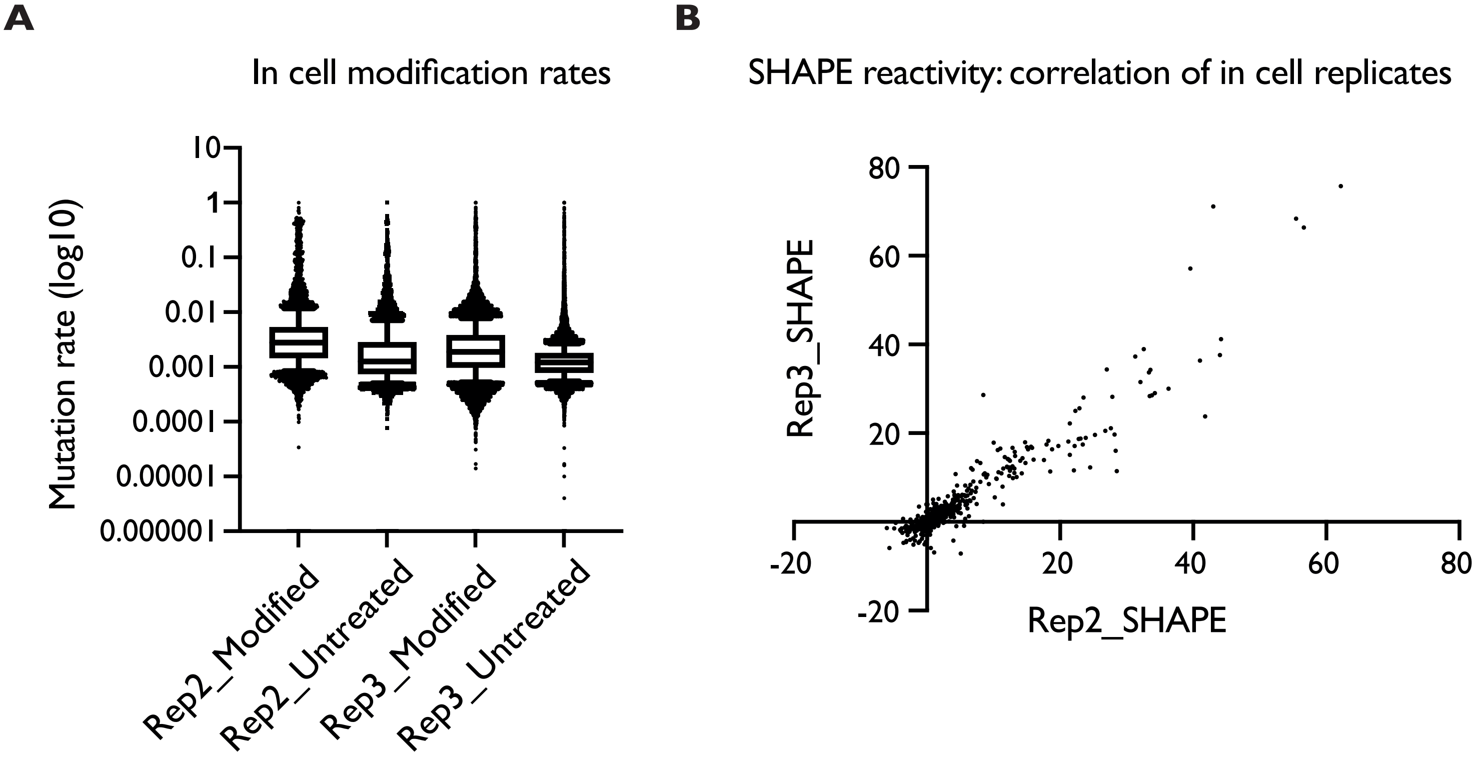


**C**

### SHAPE mapper quality control checks Replicate 2

Read depth check: 98.8% (5266/5331) nucleotides meet the minimum read depth of 5000.

Mutation rate check: 74.7% (3933/5266) nucleotides have positive mutation rates above background.

High background check:1.1% (60/5266) nucleotides have high background mutation rates. Not too many nucleotides with high background mutation rates.

Number highly reactive check: 12.5% (657/5266) nucleotides show high apparent reactivity.

### SHAPE mapper quality control checks Replicate 3

Read depth check: 94.9% (5057/5331 nucleotides meet the minimum read depth of 5000

Mutation rate check: 77.5% (3920/5057) nucleotides have positive mutation rates above background

High background check:1.8% (91/5057) nucleotides have high background mutation rates. Not too many nucleotides with high background mutation rates.

Number highly reactive check:12.8% (648/5057) nucleotides show high apparent reactivity.

**Supplemental Figure 1:** In vivo SHAPE-MaP yields sufficient quality of data to enable structure prediction and motif discovery. A: Mutation rates for two biological replicates across the entire sequence of NORAD. B: The Pearson's correlation of the two in vivo SHAPE replicates across full length NORAD is 0.9475. This is consistent with previously published Pearson's values for SHAPE reactivities of in vivo modified long non-coding RNAs (Smola et al., 2016). C: SHAPEmapper 2 Quality control statistics for the two in cell replicates,
