## Supplemental Figure 3 for "Discovery of a Well-Folded Protein Interaction Hub Within the Human Long Non-Coding RNA *NORAD*"

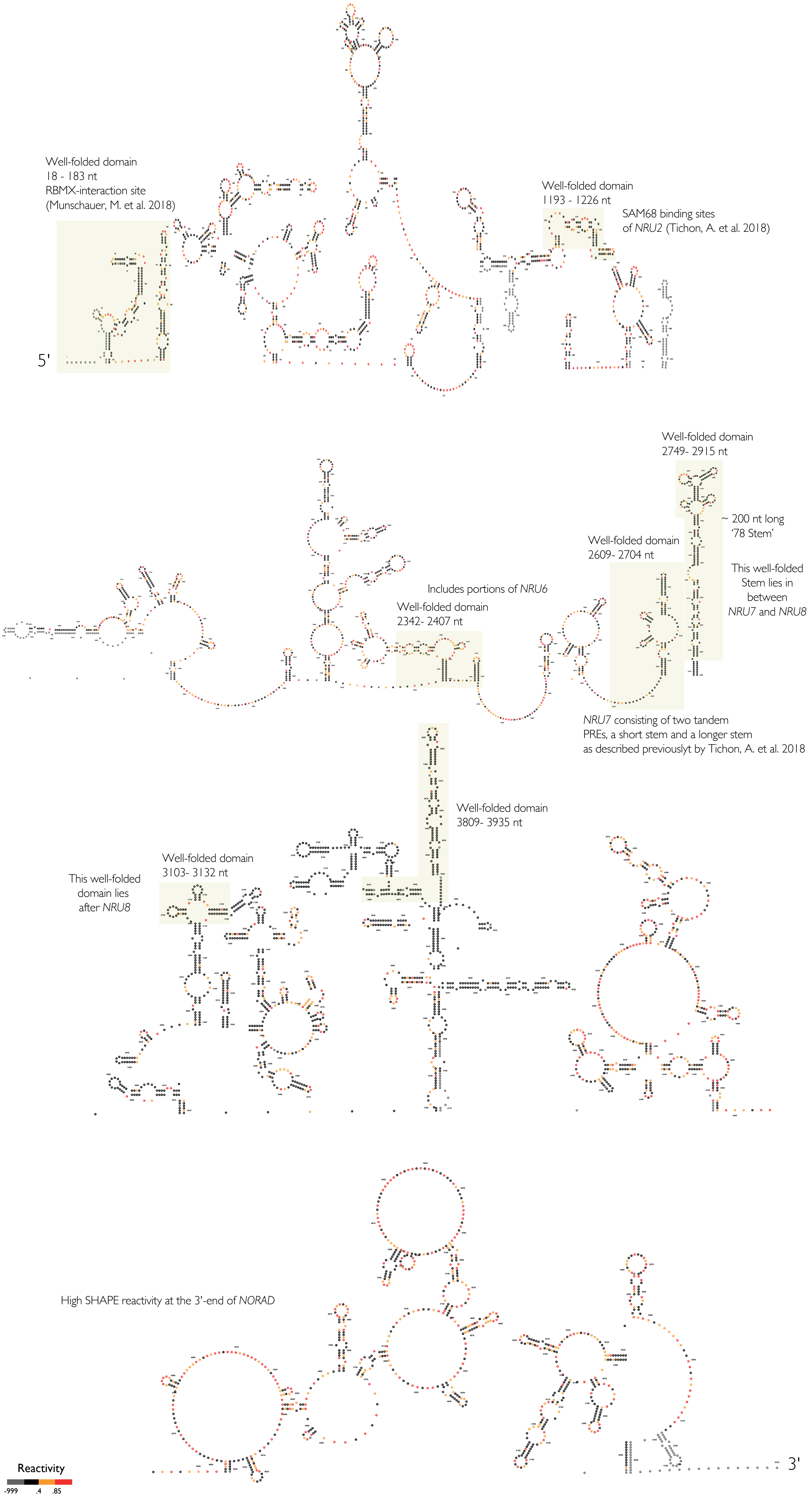


**Supplemental Figure 3:** The *in cell* secondary structural map of full-length *NORAD*. SHAPE reactivities are color coded, as shown in the legend. Yellow boxes indicates regions that are well-folded and highly structured (using the low Shannon and low SHAPE metrics). *In cell* replicate 3 data were used to generate this structural map. The structure was visualized using the program StructureEditor.
