## Supplemental Figure 4 for "Discovery of a Well-Folded Protein Interaction Hub Within the Human Long Non-Coding RNA *NORAD*"

**A**

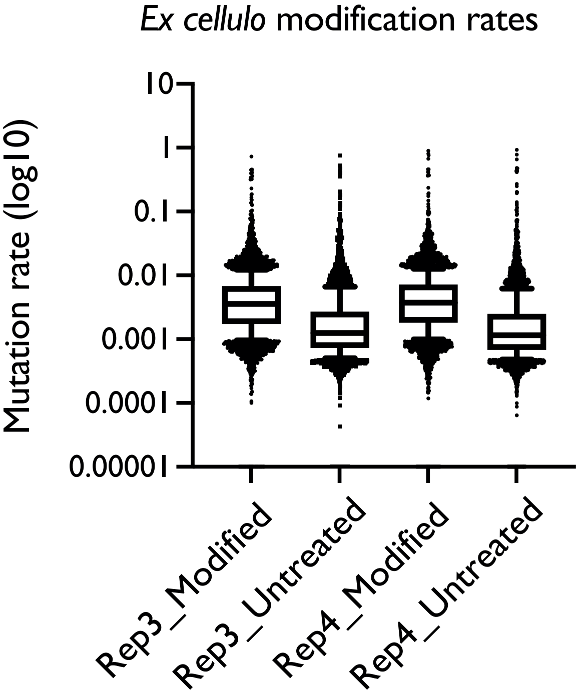


**B**

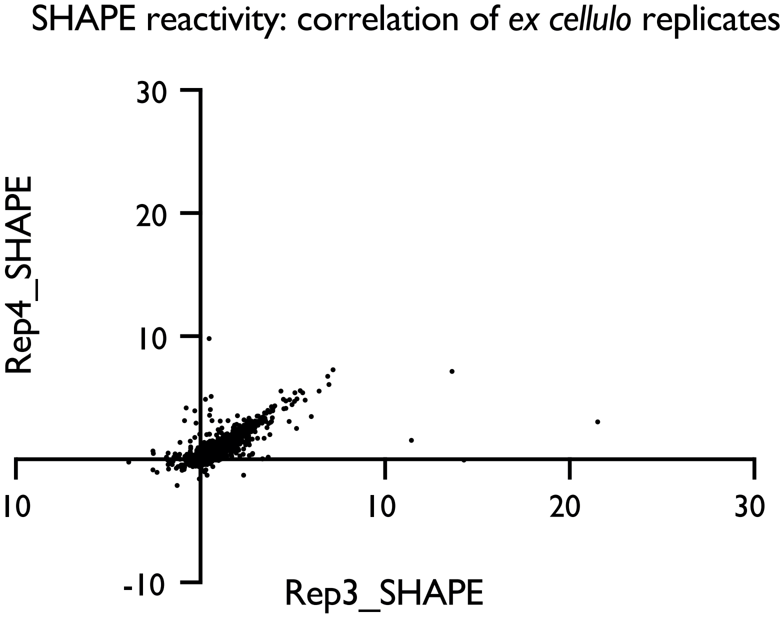


**C**

### SHAPE mapper quality control checks Replicate 3

Read depth check: 98.9% (5273/5331) nucleotides meet the minimum read depth of 5000

Mutation rate check: 84.8% (4472/5273) nucleotides have positive mutation rates above background

High background check: 0.6% (32/5273) nucleotides have high background mutation rates. Not too many nucleotides with high background mutation rates.

Number highly reactive check: 13.4% (706/5273) nucleotides show high apparent reactivity.

### SHAPE mapper quality control checks Replicate 4

Read depth check: 98.7% (5264/5331) nucleotides meet the minimum read depth of 5000

Mutation rate check: 88.5% (4659/5264) nucleotides have positive mutation rates above background

High background check: 0.6% (29/5264) nucleotides have high background mutation rates. Not too many nucleotides with high background mutation rates.

Number highly reactive check: 16.5% (871/5264) nucleotides show high apparent reactivity.

**Supplemental Figure 4:** *Ex cellulo* SHAPE-MaP yields sufficient quality of data to enable structure prediction and motif discovery. A: Mutation rates for two biological replicates across the entire sequence of *NORAD*. B: The Pearson's correlation of the two *ex cellulo* SHAPE replicates across full length *NORAD* is 0.7861. This is consistent with previously published Pearson's values for SHAPE reactivities of *in vivo* modified long non-coding RNAs (Smola et al., 2016). C: SHAPEmapper 2 Quality control statistics for the two *ex cellulo* replicates,.
