## Supplemental Figure 5 for "Discovery of a Well-Folded Protein Interaction Hub Within the Human Long Non-Coding RNA *NORAD*"

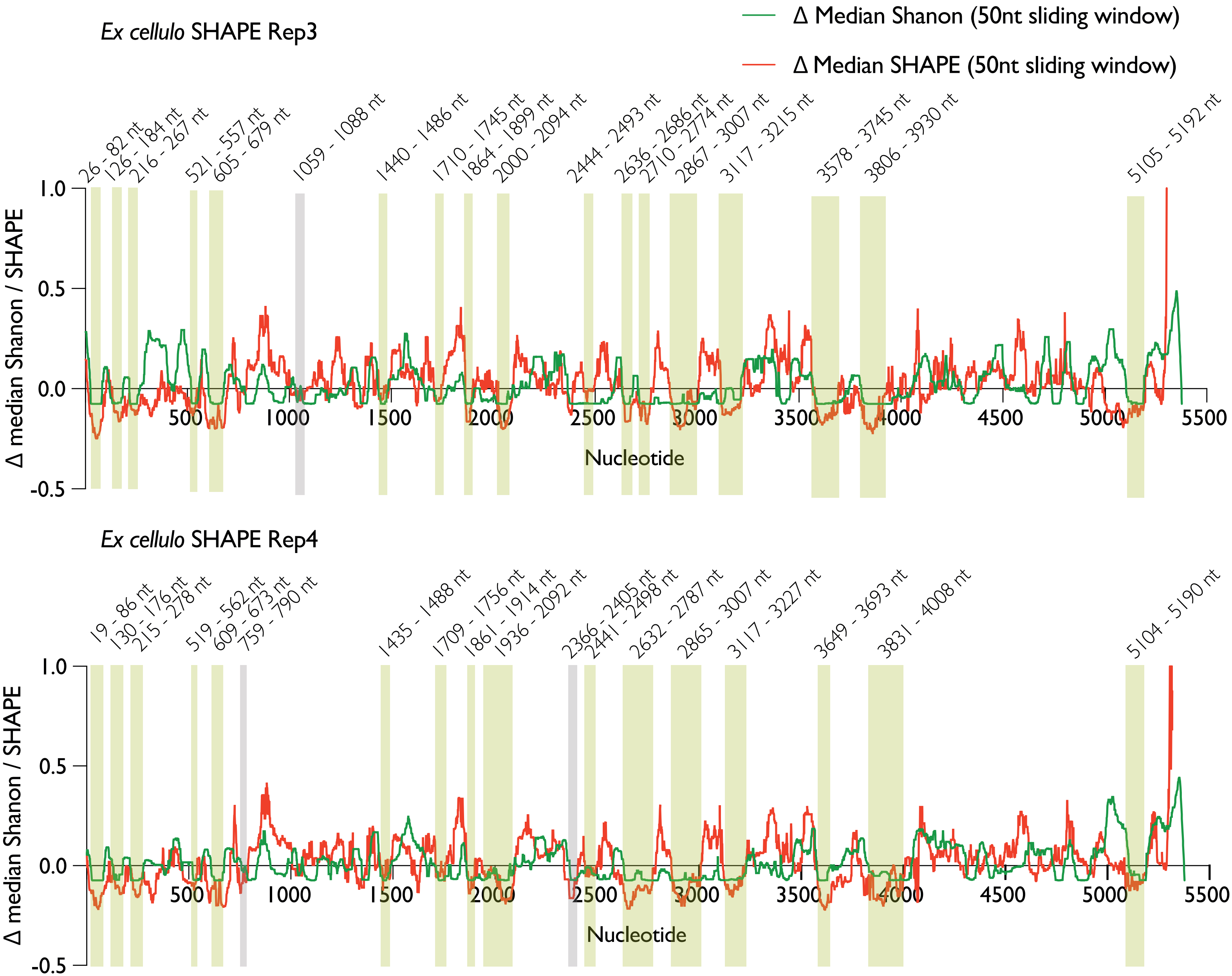


**Supplemental Figure 5:** *Ex cellulo* SHAPE-MaP of *NORAD* reveals several well-folded structural motifs. Comparison of local median Shannon entropy and SHAPE reactivities calculated from *ex cellulo* SHAPE-MaP experiment across the two replicates. Nucleotide numbers are indicated on X-axis. Yellow boxes represent well-folded motifs found in both the replicates. Grey boxes represent well-folded domains found in only one of the two replicates.
