## Supplemental Figure 6 for "Discovery of a Well-Folded Protein Interaction Hub Within the Human Long Non-Coding RNA *NORAD*"

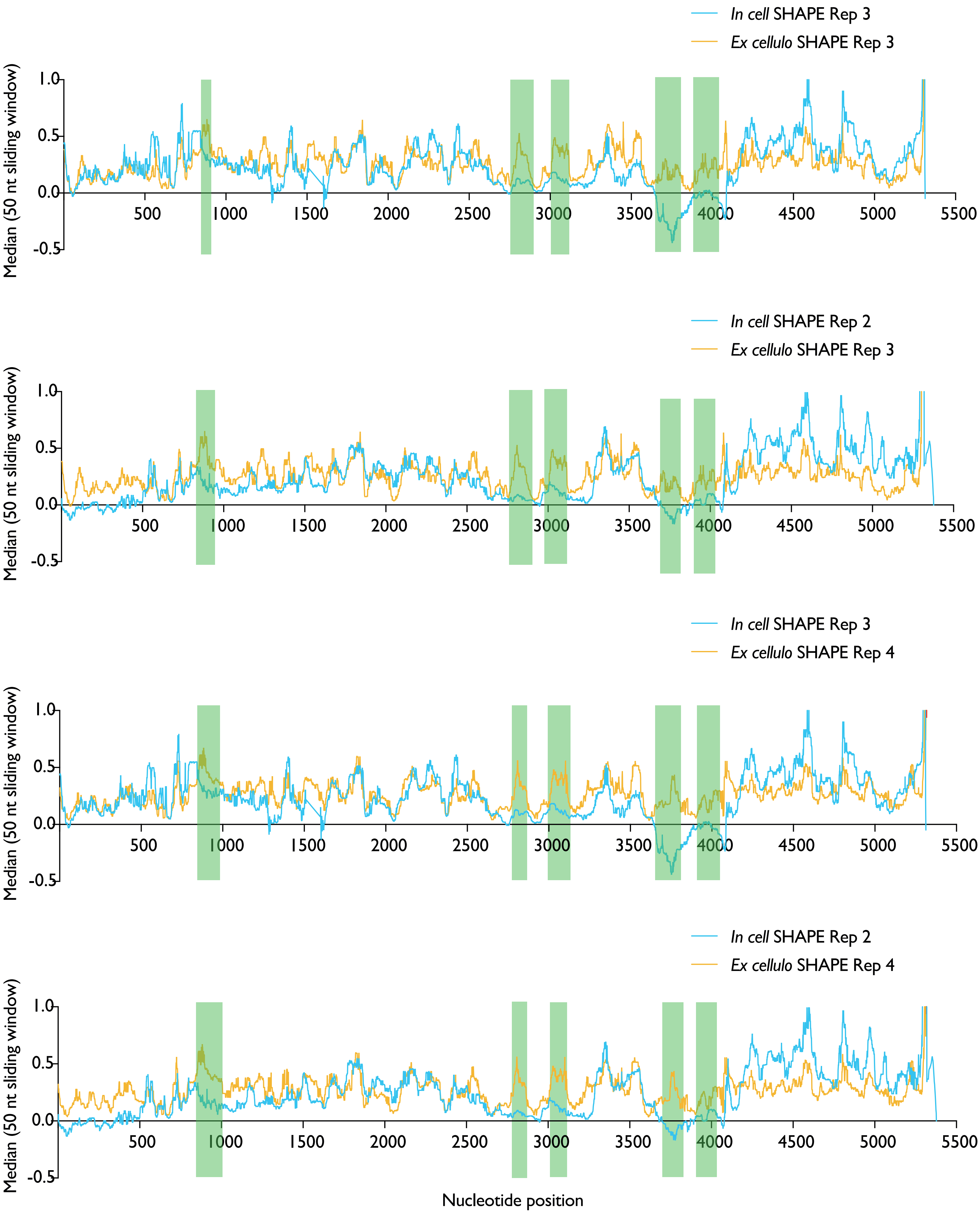


**Supplemental Figure 6 :** Identification of protein binding hubs within *NORAD* using *ex cellulo* SHAPE-MaP. Comparison of *in cell* and *ex cellulo* SHAPE reactivities in 50-nt sliding windows across all the replicates. Green boxes indicates regions protected by cellular environment, In total, five protein binding hubs are found across all the four compared samples.
