## Supplemental Figure 7 for "Discovery of a Well-Folded Protein Interaction Hub Within the Human Long Non-Coding RNA *NORAD*"

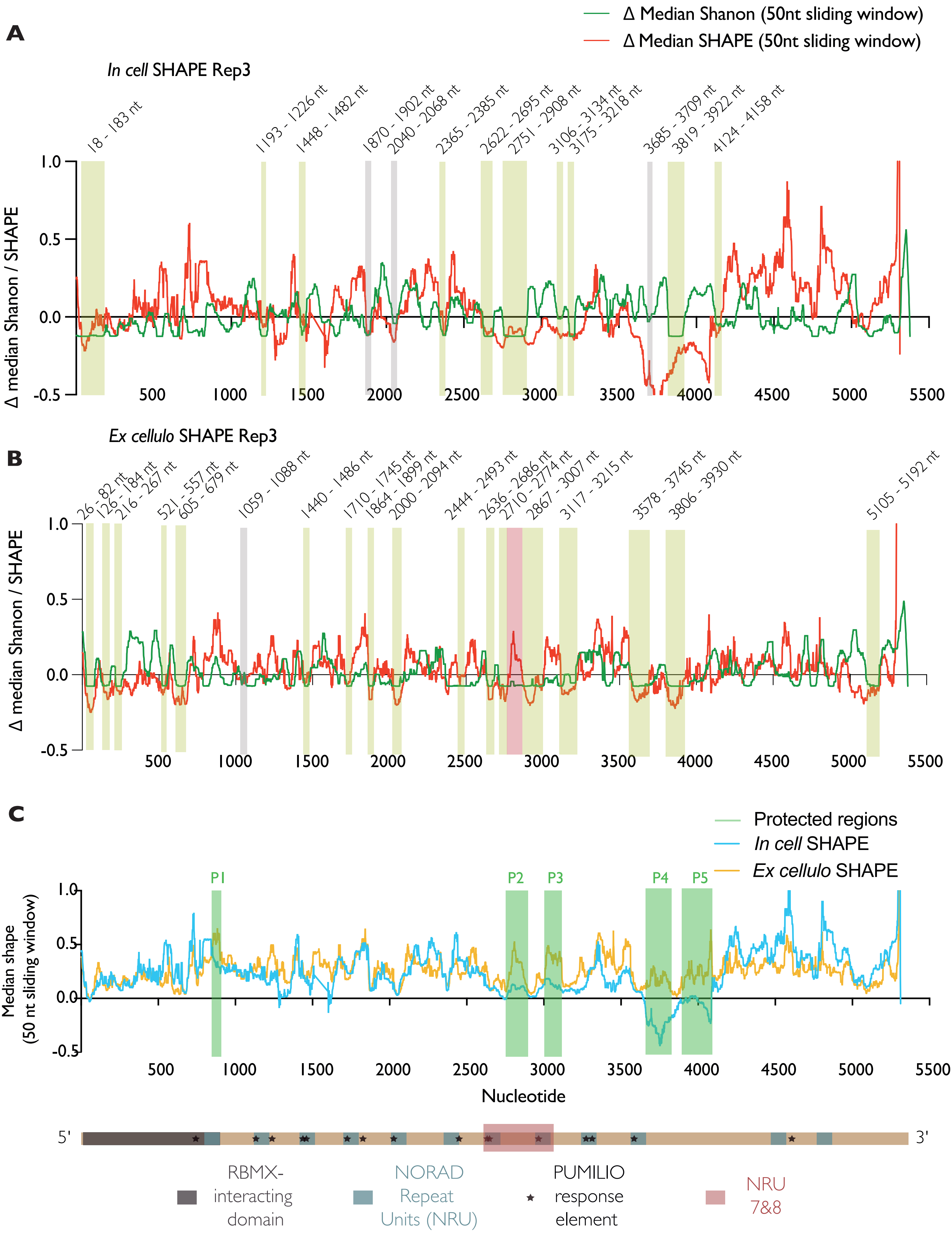


**Supplemental Figure 7 :** Comparison of the low-SHAPE and low-Shannon (lowSS) regions between *in cell* and *ex cellulo* SHAPE-MaP of *NORAD* reveals sites that becomes ordered likely upon protein binding. Yellow boxes represent well-folded motifs found in both the conditions. Grey boxes represent well-folded domains found in only one of the *in cell* or *ex cellulo* replicates. Red box represent a region between 2775 - 2865 nt of *NORAD* that appears to be ordered and well-folded in the *in cell* condition, whereas becomes reactive upon depletion of proteins.
