## Supplemental Figure 8 for "Discovery of a Well-Folded Protein Interaction Hub Within the Human Long Non-Coding RNA *NORAD*"

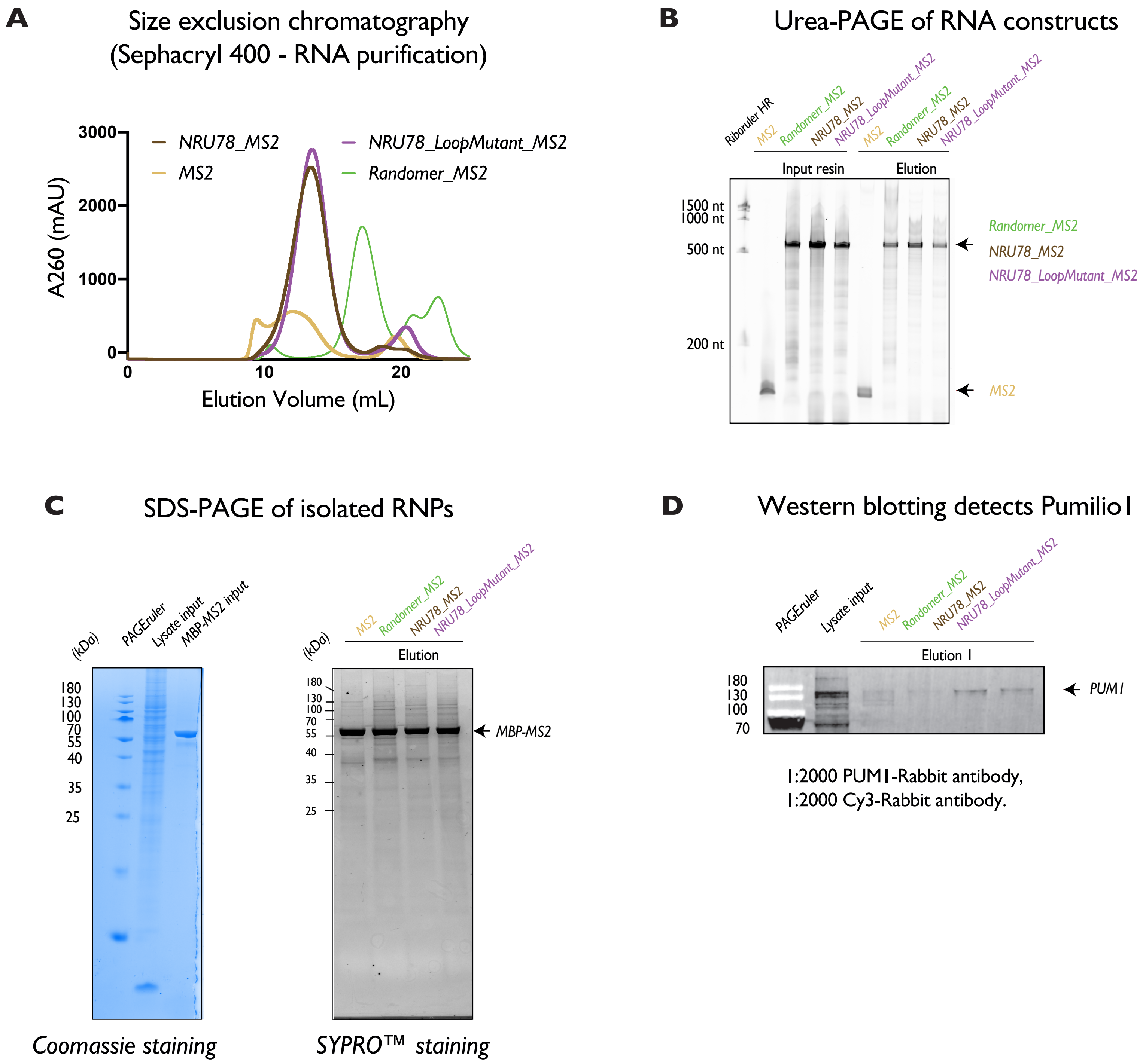


**Supplemental Figure 8:** *In vitro* pull-downs coupled with quantitative mass spectrometry identified protein partners of *NORAD* A: Native purification of RNA constructs used in the pull-down experiments. Size exclusion chromatogram of *in vitro* transcribed RNAs using Sephacryl 400 column in 50 mM HEPES pH 7, 150 mM KCL, 0.5mM EDTA buffer and run at 0.5 ml/min flow rate. B: Visualizing the purity and size of the RNA constructs used in the pull-downs (input), and the RNA fractions eluted after the pull downs (Elution). The samples were run on a 6% polyacrylamide-7M Urea gels. C: Visualizing the input lysate, MS2-MBP bait protein used in the pull-downs by NuPAGE™ 4 to 12%, Bis-Tris PAGE (ThermoFisher) and coomassie staining (left). The protein content and composition of the RNPs eluted in the pull-dows are visualized by SYPRO™ staining (ThermoFisher) (right). D: Western Blot analysis on the eluted RNPs to confirm the specificity of the pull-downs. Pumilio 1 polyclonal anti-rabbit antibody (Protein Tech) were used in the experiment. 7 uL of elution samples and ~ 4 uL of lysate samples were loaded per well and run on a 14-12% NuPAGE commercial gels at 80V for1 hour, before blotting for Pumilio1.
