## Supplemental Figure 9 for "Discovery of a Well-Folded Protein Interaction Hub Within the Human Long Non-Coding RNA *NORAD*"

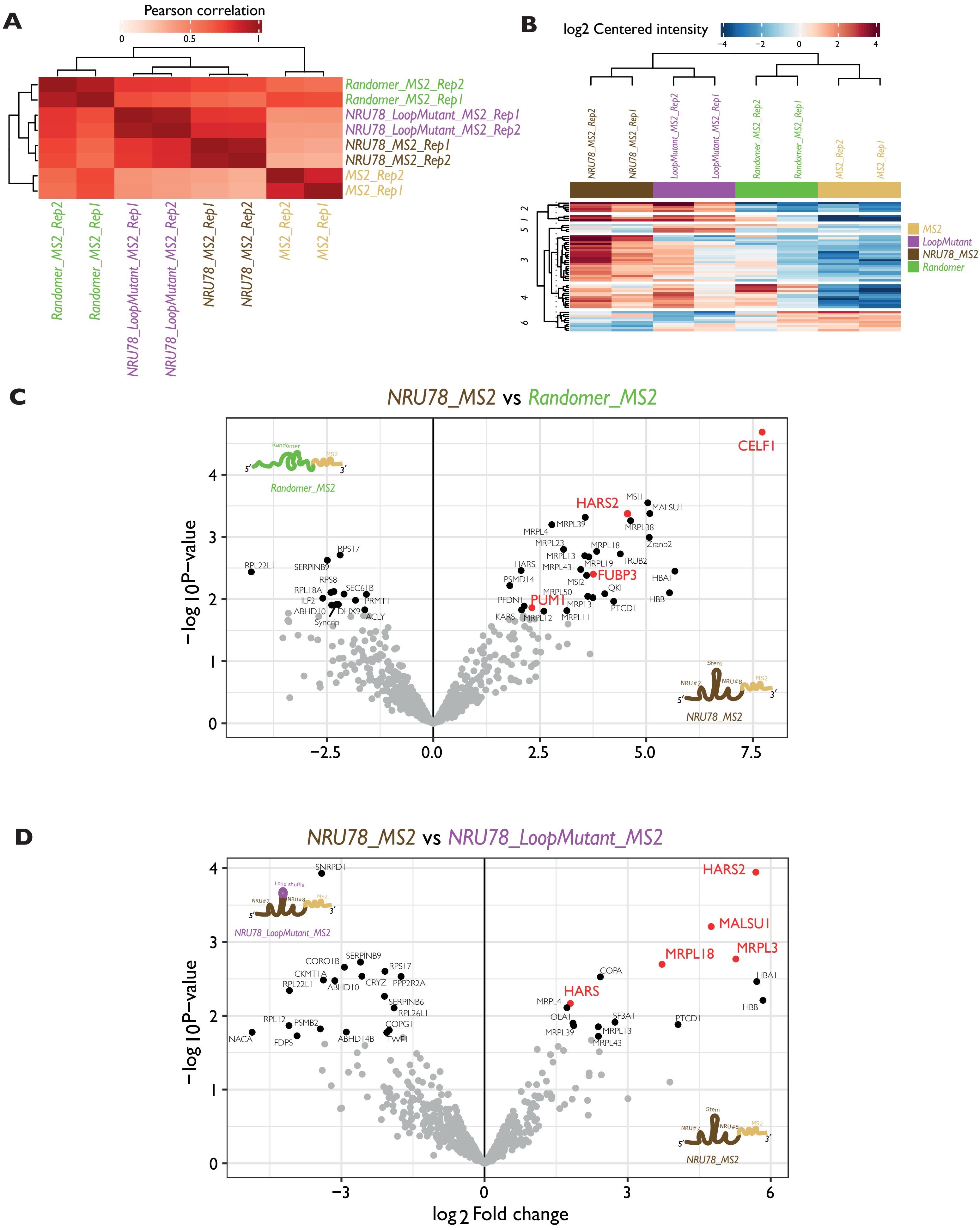


**Supplemental Figure 9:** *In vitro* pull-downs coupled with quantitative mass spectrometry identified protein partners of *NORAD* A: Pearson's correlation martix between the different samples used in the pull-downs. B: Heatmap representation providing an overview of all significant proteins (rows) in all replicates of all the four RNA constructs used in the pull-downs. C: Volcano plots showing specific enrichment of proteins in *NORAD NRU78* pull-downs in comparison with the randomer RNA control pull-downs. D: Volcano plots showing specific enrichment of proteins in *NORAD NRU78* pull-downs in comparison with the *NRU78* RNA with its *78* stem loop mutated control pull-downs.
