## Supplemental Figure 10 for "Discovery of a Well-Folded Protein Interaction Hub Within the Human Long Non-Coding RNA *NORAD*"

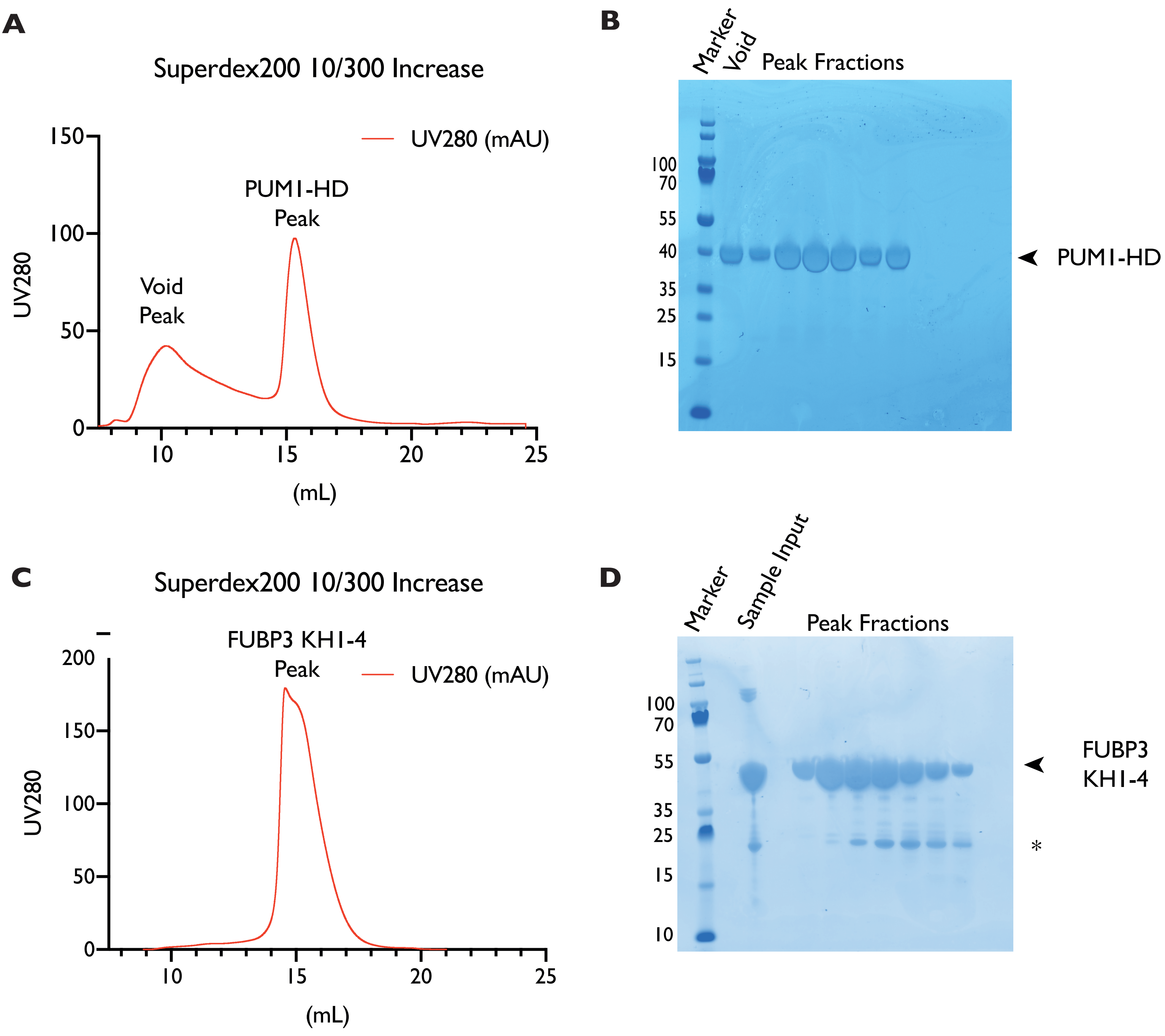


**Supplemental Figure 10:** Reconstitution of a minimal *NORAD*-Pumilio1-FUBP3 complex using purified components. A: Size exclusion chromatogram of PUM1-HD protein shows a homogeneous and well-dispersed peak. B: The corresponding SDS-PAGE of the peak fractions from A reveals pure PUM1-HD. All the peak fractions were pooled together, concentrated to 70 μM and stored at -80 °C for future biochemical studies. C: Size exclusion chromatogram of FUBP1 KH-14 protein shows a homogeneous and well-dispersed peak (with a shoulder). D: The corresponding SDS-PAGE of peak fractions from C reveals pure FUBP3 KH-14. * represents possible degradation and explains the right-shoulder of the peak. The first three peak fractions were pooled together, concentrated to 259 μM and stored in -80 °C for future biochemical experiments.
